# Conformation governs cellular behaviors of LRRK2 Parkinson’s disease variants

**DOI:** 10.64898/2026.10.30.763152

**Authors:** Kathryn S. Hatch, Katherine J. Surridge, Wei Gai, Brookelyn Quan, Gaurav Kumar, Nicholas Lattal, Robert Abrisch, Niels Pagani, Andres E. Leschziner, Samara L. Reck-Peterson

## Abstract

Gain-of-function of leucine-rich repeat kinase 2 (LRRK2) is a common cause of Parkinson’s disease (PD). In its autoinhibited state, the leucine-rich repeats of LRRK2 block the kinase active site, preventing substrate phosphorylation. Displacing the repeats (‘unlatching’) primes LRRK2 for activity. Preferential unlatching causes increased substrate phosphorylation of the PD allele LRRK2[R1441H], but not LRRK2[G2019S]. Here, we define a large class of PD-associated ‘unlatching’ mutations based on their effect on LRRK2 conformation. Unlatching mutants show increased localization to membranes, the site of LRRK2’s Rab substrates. Unlatching mutants are also characterized by decreased LRRK2 S935 phosphorylation, which we define as a biomarker of conformation. This framework resolves the mechanism of variants such as I2020T, which structural studies could not predict. Overall, we establish unlatching as a unifying mechanism for most PD-linked variants and describe measurable readouts for identifying LRRK2-driven disease.

## Introduction

Parkinson’s disease (PD) is the second most prevalent neurodegenerative disease worldwide^1,2^. Mutations in leucine-rich repeat kinase 2 (*LRRK2*) are one of the most common causes of genetic PD^3,4^. All established PD-linked mutations are gain-of-function variants that increase LRRK2mediated phosphorylation of Rab GTPase substrates on organelle membranes^5–7^. Dysregulation of LRRK2 kinase activity has also been implicated in idiopathic PD, positioning LRRK2 as a potential major therapeutic target^8–10^. However, the molecular mechanisms driving increased Rab substrate phosphorylation by PD-associated LRRK2 variants are poorly understood. Defining how gain-of-function is conferred by PD-linked mutations is critical for understanding disease mechanisms and informing the development of targeted therapeutics.

LRRK2’s C-terminal half contains its two enzymatic modules, a Ras of Complex (ROC) GTPase domain and a serine/threonine kinase^11^. The GTPase and kinase are connected by a flexible C-terminus Of ROC (COR) scaffolding domain, and followed by a WD40 domain^11^. Collectively, these 4 modules are named ‘RCKW’, for **R**OC, **C**OR, **K**inase and **W**D40 (Figure 1A). LRRK2’s N-terminal half is comprised of three repeat domains: Armadillo (ARM), Ankyrin (ANK), and Leucine-Rich Repeats (LRR)^12^. The position of these repeat domains plays a key role in regulating LRRK2 substrate phosphorylation. In the autoinhibited state, the leucine-rich repeats are positioned across the kinase, physically blocking substrate access to the active site and constraining the kinase in its inactive state^12^. Unlatching of the leucine-rich repeats is required for (and upstream of) substrate phosphorylation (Figure 1B). While PD-associated mutations in LRRK2 have been identified across the entire protein, most are found in the catalytic C-terminal half (Figure 1A)^13^. Two widely studied pathogenic mutations are found in LRRK2’s kinase: a glycine to serine substitution at residue 2019 (G2019S) and an isoleucine to threonine substitution at residue 2020 (I2020T)^14–17^. Arginine 1441 (R1441) in the ROC domain is the second-most frequently mutated residue associated with LRRK2-driven PD after G2019, with three pathogenic substitutions reported: R1441H, R1441C, and R1441G (LRRK2[R1441H/C/G])^18–20^.

**Figure 1.**
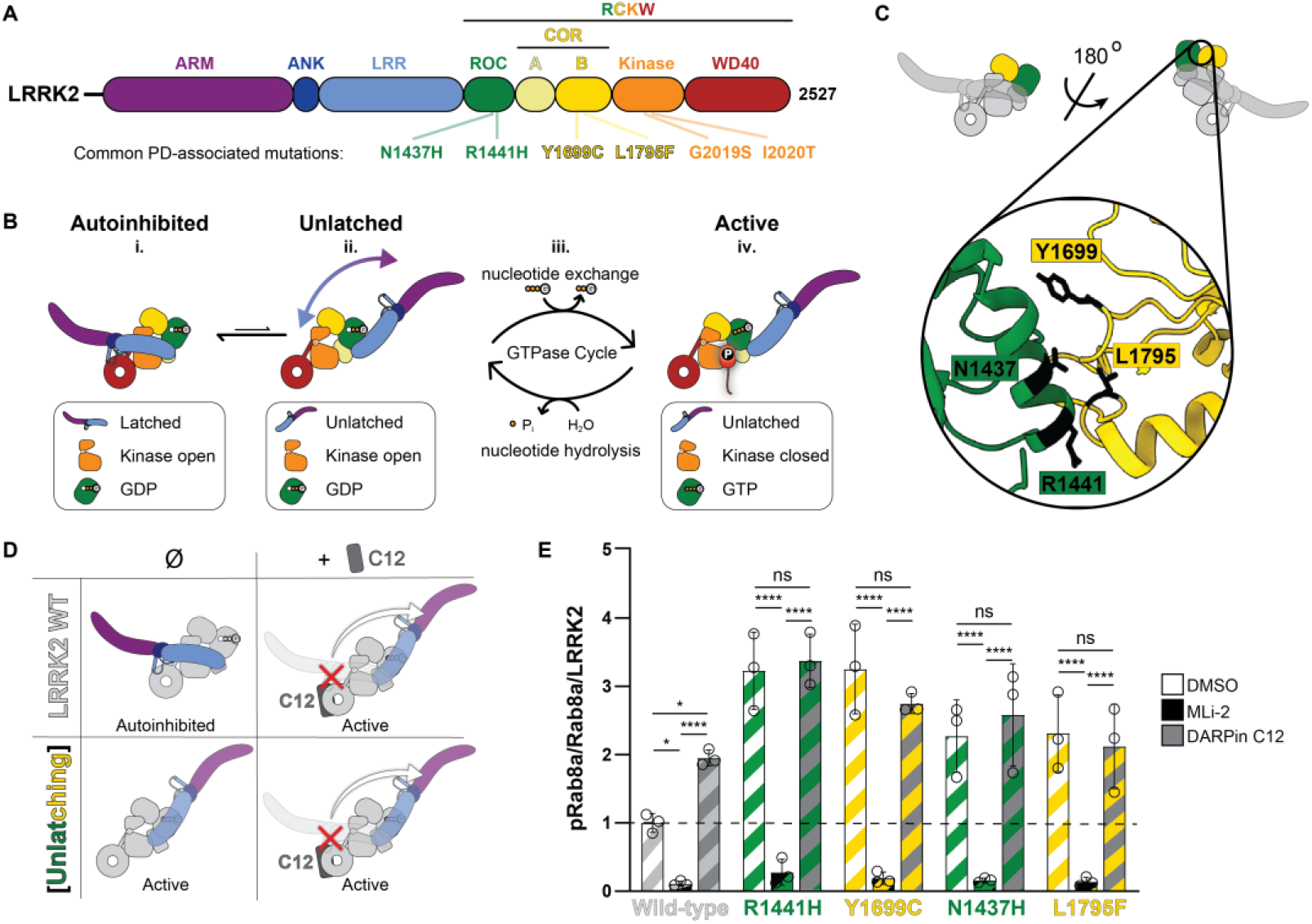
Mutations along the ROC:COR-B interface favor the unlatched conformation of LRRK2. **(A)** Schematic representation of the domain organization of LRRK2. LRRK2 is composed of three N-terminal repeat domains (Armadillo; ARM, Ankyrin; ANK and Leucine-Rich Repeats; LRR), a GTPase domain (Roc Of Complex; ROC), a linker domain (C-terminal Of Roc; COR), a serine/threonine kinase and a C-terminal WD40 domain. The four C-terminal modules are collectively named ‘RCKW’ for ROC, COR, Kinase and WD40. Some of the most common PD-associated mutations (and those described in this study) are annotated. This color scheme for the domains will be used throughout. **(B)** Cartoon representation of the current model for LRRK2’s activation cycle. In cells, LRRK2 primarily exists in the cytoplasm in an autoinhibited state, where the leucine-rich repeats are latched across the open, inactive kinase domain and the ROC domain is bound to GDP (i). In a small population of LRRK2, the repeats transiently unlatch from their position across the kinase active site (ii). Unlatching the repeats relieves the conformational restraint across the ROC-COR module, lowering the affinity for GDP. As GTP is more abundant in the cell, this allows for nucleotide exchange to GTP (iii). GTP-binding stabilizes the active conformation, with the repeats unlatched and the kinase in a closed, active state. This stably active state is thought to be responsible for the majority of substrate phosphorylation (iv). GTP must be hydrolyzed to GDP for the leucine-rich repeats to stably relatch across the kinase again (iii). **(C)** Close association between the ROC and COR-B domains is a key feature of the auto-inhibited state (PDB: 9OXH^23^). The ROC:COR-B interface is a hotspot for PD-associated gain-of-function mutations that are predicted to weaken the stability of this interface. **(D)** Predicted predominant conformational state of LRRK2[WT] and ‘unlatching mutants’ in the absence or presence of DARPin C12. **(E)** Quantification of phosphorylated Rab8a (pT72)/total Rab8a/total LRRK2, normalized to the LRRK2 [WT] DMSO condition from 3 (n=3) independent western blots. 293T cells overexpressing GFP-tagged Rab8a and the indicated GFP11-tagged LRRK2 variants with or without FLAG-tagged DARPin C12 were treated with either DMSO or the LRRK2 kinase inhibitor MLi-2 (5 μM, 1 hour). Blots were processed using ImageJ and the intensity of protein bands were analyzed using a two-way ANOVA with Tukey’s multi-comparison correction. * p≤0.05; **** p≤0.0001; ns=non-significant. Error bars +/- SD. Representative immunoblots of this analysis can be found in Supplementary Figure 1B and C.

Other common PD-associated mutations include N1437H/D in the ROC domain and Y1699C and L1795F in the COR-B domain (Figure 1C); the presence and prevalence of multiple gain-of-function mutations in the ROC GTPase and the adjacent COR-B domain suggests potential allosteric regulation of LRRK2 activity^21,22^.

Recently, we uncovered the intramolecular mechanism regulating LRRK2 activation^23^ (Figure 1B). In the autoinhibited conformation, the latching of the N-terminal repeats physically constrains the kinase in an inactive state, rigidifying the ROC:COR-B interface, and favoring GDP binding at the ROC domain^23^. Unlatching of the leucine-rich repeats (the ‘unlatched’ state) allows for conformational changes at the ROC:COR-B interface and nucleotide exchange to GTP in the ROC domain^23^. In the GTP-bound ‘active’ state, the leucine-rich repeats are sterically prohibited from re-latching across the kinase; this conformation is likely responsible for most substrate phosphorylation (Figure 1B). This mechanistically explains why pathogenic mutations at R1441 are gain-of-function. R1441 in the ROC domain stabilizes the autoinhibited state through interactions with the COR-B domain, and PD-associated mutations at R1441 destabilize this interface, favoring unlatching and priming LRRK2 for kinase activity (Figure 1D)^23^. This mode of action is mechanistically distinct from that of G2019S, which is hypothesized to act directly on the kinase reaction. These insights raise three questions: is preferential unlatching of the leucine-rich repeats a common mechanism of action for other PD-associated LRRK2 variants; does this mechanism form the molecular basis behind reported R1441 mutant-specific behaviors^7,24–27^; and does this mechanism account for the cellular behaviors of other PD-associated variants as well?

R1441 mutant LRRK2 exhibits distinct molecular and cellular behaviors compared to both wild-type LRRK2 and the most common PD-associated mutant G2019S. The basis for these differences is not well understood^28,29^. While both G2019S and R1441 mutants are gain-of-function variants, the extent of their effects on Rab substrate phosphorylation and LRRK2 phosphorylation state differs^7,26,30,31^. Phosphorylation of LRRK2 by upstream kinases at S935 is reduced in R1441 mutant LRRK2 compared to either wild-type or G2019S variants^7,30,31^. In contrast, autophosphorylation of LRRK2 at S1292 is increased in both variants, albeit to differing extents^7,26^. Mutations at R1441 also reduce GTPase activity and increase GTP binding relative to both wild-type and G2019S mutant LRRK2^32–34^. Additionally, under certain conditions R1441 mutant LRRK2 exhibits increased recruitment to intracellular structures, including microtubules and membranes, where its Rab substrates are located^24,25,35,36-39^. The molecular mechanisms underlying the distinct behaviors associated with R1441 mutant LRRK2 are unresolved.

In this study, we establish unlatching of the leucine-rich repeats as a common mechanism underlying many PD-associated LRRK2 mutations, while highlighting LRRK2 mutant G2019S as a mechanistically distinct case. Using complementary biochemical and fluorescent imaging approaches, we show that this mechanism of action has cellular consequences that extend beyond substrate phosphorylation to membrane localization, LRRK2 phosphorylation profile, and nucleotide state at the ROC GTPase domain. We define a class of PD-associated ‘unlatching’ mutations based on their effect on LRRK2 conformation, which correlates with increased membrane localization and LRRK2 S935 dephosphorylation. These mechanistic distinctions may be significant for understanding the phenotypic heterogeneity seen in LRRK2-associated familial PD, and for understanding the etiology of LRRK2-driven PD where LRRK2 is not genetically implicated.

## Results

### PD-associated mutations at the ROC:COR-B interface facilitate repeat unlatching

The ROC:COR-B interface is a hotspot for PD-associated gain-of-function mutations (Figure 1A,C)^13^. We hypothesized that all mutations at this interface would increase Rab substrate phosphorylation by promoting unlatching of the leucine-rich repeats, as we previously showed for LRRK2[R1441H] (Figure 1D)^23^. In our earlier work, we identified Designed Ankyrin Repeat Proteins (DARPins) that bound to LRRK2’s WD40 domain. We found that one of these, DARPin C12, leads to increased Rab phosphorylation by favoring the unlatched state of LRRK2 (Figure 1D; Supplementary Figure 1A)^37^. The presence of DARPin C12 in cell-based kinase assays led to increased Rab8a phosphorylation by LRRK2[WT] and LRRK2[G2019S], but did not affect substrate phosphorylation by LRRK2[R1441H], indicating that LRRK2[R1441H] intrinsically favors the unlatched conformation^23^. To determine if other mutations at the ROC:COR-B interface behave similarly, we measured Rab8a phosphorylation in 293T cells co-expressing Rab8a, DARPin C12 and either LRRK2[WT], LRRK2[R1441H], LRRK2[N1437H], LRRK2[Y1699C] or LRRK2[L1795F]. We observed no increase in Rab8a phosphorylation in the presence of DARPin C12 in cells expressing these ROC:CORB interface mutations (Figure 1E; Supplementary Figure 1B,C). These data suggest that mutations on either side of the ROC:COR-B interface can be mechanistically classified as ‘unlatching mutants’ that increase Rab phosphorylation by facilitating the unlatching of the leucine-rich repeats, which primes LRRK2 for kinase activity.

### Phosphorylation profiles of PD-associated LRRK2 variants are driven by conformation

LRRK2 is phosphorylated by upstream kinases (including Protein Kinase A^40^, Casein Kinase 1^41^, and IκB family kinases^42^) at biomarker sites (S910 and S935) in a flexible loop region between the ANK and LRR domains (Figure 2A)^43,44^. These sites are dephosphorylated by the phosphatases PP1^45^ and PP2A^46^ following inhibition of LRRK2 by type I kinase inhibitors^47^; dephosphorylation at S935 is therefore used as a readout of target engagement and activity in both pre-clinical studies and clinical trials targeting LRRK2^48,49^. Canonical engagement of type I inhibitors requires LRRK2 to be in its active-like, closed kinase state, consistent with the unlatched conformation (Supplementary Figure 1A)^50^. In contrast, treatment with type II kinase inhibitors that engage LRRK2 in the kinase open, autoinhibited state, maintains phosphorylation at these biomarker sites^47,50,51^.

**Figure 2.**
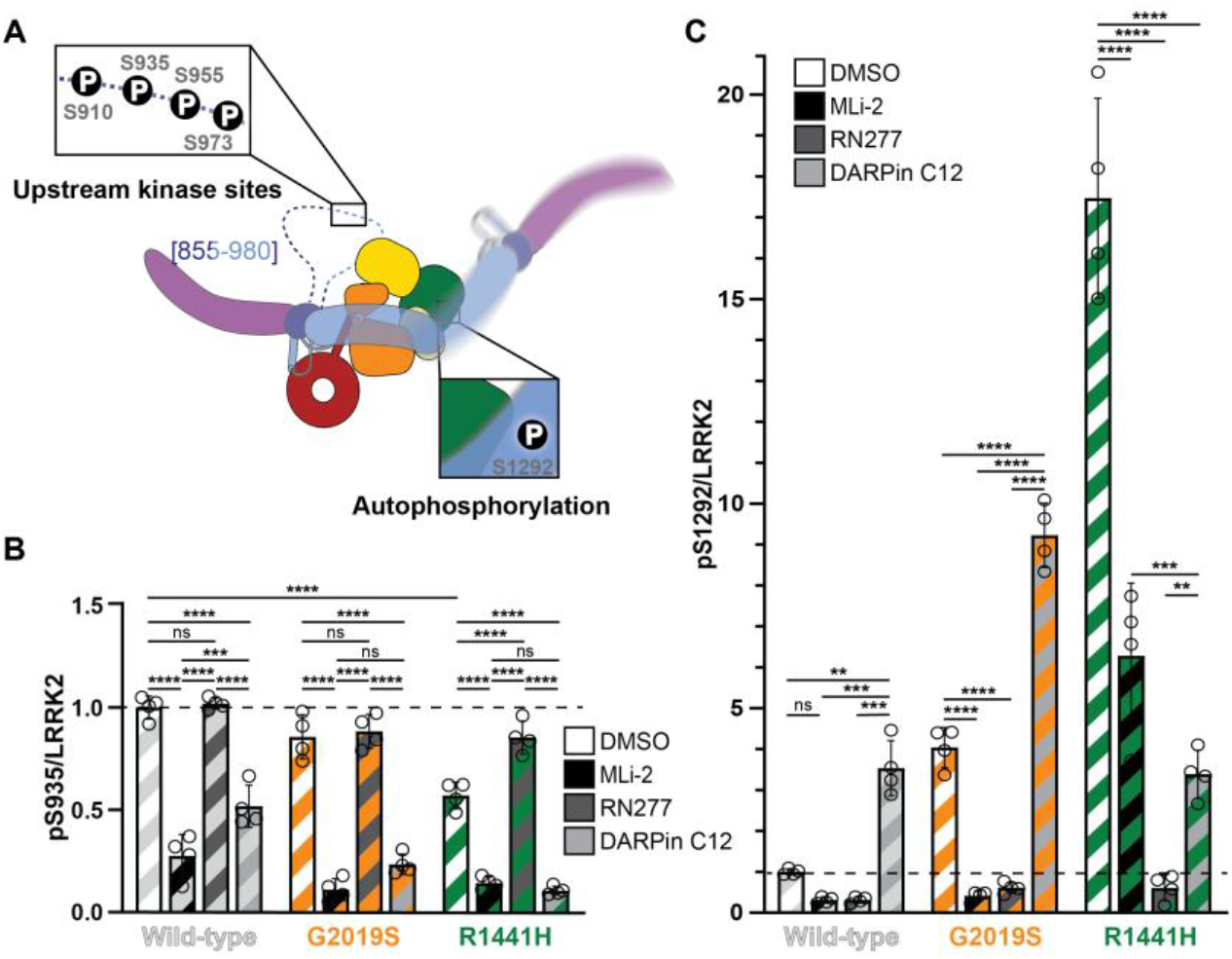
Conformation drives distinct phosphorylation profiles of LRRK2. **(A)** Cartoon depiction of LRRK2 with the phosphorylation sites relevant to this study annotated. LRRK2 is phosphorylated by upstream kinases at multiple serines in a disordered loop region (residues 855-980) between the ANK and LRR domains and undergoes autophosphorylation at serine 1292 in the LRR domain. **(B)** Quantification of phosphorylated LRRK2 (pS935)/total LRRK2, normalized to the LRRK2[WT] DMSO condition from 4 (n=4) independent western blots. 293T cells overexpressing GFP11-tagged LRRK2 variants with or without FLAG-tagged DARPin C12 were treated with DMSO, MLi-2 (1 μM) or RN277 (10 μM) for 3 hours. Data were processed as previously described using a two-way ANOVA corrected for multiple comparisons using Tukey’s test. *** p≤0.001; **** p≤0.0001; ns=non-significant. Error bars +/- SD. Representative immunoblots related to this analysis can be found in Supplementary Figure 2A. **(C)** Quantification of phosphorylated LRRK2 (pS1292)/total LRRK2, normalized to the LRRK2[WT] DMSO condition from 4 (n=4) independent western blots. Samples are the same as those for Figure 2B, rerun under identical conditions and analyzed as previously described. ** p≤0.01; *** p≤0.001; **** p≤0.0001; ns=non-significant. Error bars +/- SD. Representative immunoblots related to this analysis can be found in Supplementary Figure 2B.

We hypothesized that unlatching of LRRK2’s leucine-rich repeats induced by DARPin C12 would promote dephosphorylation of S935, indicative of a conformation-specific effect. To test this, we transfected 293T cells with either LRRK2[WT], LRRK2[G2019S] or LRRK2[R1441H] and co-transfected with DARPin C12 or treated with the type I inhibitor MLi-2 or the type II inhibitor RN277^47^. We then measured relative S935 phosphorylation levels using a phospho-S935 specific antibody (Supplementary Table 1). We found that LRRK2[WT], LRRK2[G2019S] and LRRK2[R1441H] all showed decreased phosphorylation of S935 in the presence of DARPin C12 or MLi-2 (Figure 2B; Supplementary Figure 2A). In addition, DMSO treated LRRK2[R1441H] alone showed decreased phosphorylation at S935 compared to LRRK2[WT] or LRRK2[G2019S], as has been previously reported (Figure 2B)^7,43,52^. These data support the hypothesis that reduced phosphorylation at S935 is a readout of the unlatched conformation of the repeats, favored by the ROC:COR-B destabilizing mutant R1441H and the binding mode of both type I inhibitors and DARPin C12. The extent of dephosphorylation induced by either DARPin C12 or MLi-2 was greater than LRRK2[R1441H] alone, indicating that the mutation produces only a partial shift in the conformational equilibrium, whereas excess MLi-2 or DARPin C12 drive a greater accumulation of unlatched LRRK2. Phosphorylation at S935 was maintained following treatment with the type II inhibitor RN277 in LRRK2[WT] and LRRK2[G2019S] and increased to wild type levels in LRRK2[R1441H], consistent with previous observations from structural studies^23^ that LRRK2[R1441H] retains the ability to access the autoinhibited conformation associated with S935 phosphorylation (Figure 2B). These data separate two readouts that have often been conflated; S935 phosphorylation is a biomarker for LRRK2’s conformational state, and independent from kinase activity.

In addition to phosphorylation by upstream kinases, LRRK2 is also autophosphorylated^26,53^. We next tested the effect of DARPin C12 on autophosphorylation at site S1292. Co-expression of DARPin C12 with either LRRK2[WT] or LRRK2[G2019S] increased phosphorylation at S1292, as anticipated for a condition that promotes the unlatched, active conformation (Figure 2C; Supplementary Figure 2B). Surprisingly, while LRRK2[R1441H] alone showed more phosphorylation at S1292 than either LRRK2[WT] or LRRK2[G2019S], co-expression of DARPin C12 decreased autophosphorylation of LRRK2[R1441H] at S1292. The COR-B domain mutant LRRK2[Y1699C], which shares the same mechanism of action (Figure 1E), also showed decreased autophosphorylation at S1292 in the presence of DARPin C12, indicating a common characteristic of ‘unlatching mutations’ under these conditions (Supplementary Figure 3).

### Unlatched LRRK2 preferentially localizes to membranes

LRRK2 is primarily cytosolic in cells, but must be recruited to membranes to phosphorylate its Rab GTPase substrates^54^. Several conditions have been shown to promote LRRK2 recruitment to membranes, including lysomotropic agents^55^, the VPS34 inhibitor SAR405^24^, overexpression of recruitment factors such as Rab29^25^, cytoplasmic DNA^56^, and endosome/phagosome membrane damage by intracellular pathogens^57^. PD-associated ROC:COR-B interface mutants were shown to be preferentially localized to Rab5a-positive endosomes in response to the VPS34 inhibitor SAR405 when compared to LRRK2[WT] and LRRK2[G2019S]^24^. In light of our current data (Figure 1E), we hypothesized that this may be a conformation regulated phenotype; the unlatched conformation of LRRK2 may preferentially localize to endosomes in response to SAR405 treatment. To test this, we co-expressed LRRK2[WT], LRRK2[G2019S] or LRRK2[R1441H] with or without DARPin C12 in 293T cells and treated with SAR405. We then used spinning-disk confocal microscopy and colocalization analysis to quantify LRRK2’s colocalization with Rab5a-positive endosomes. Cells expressing LRRK2[WT] did not show a significant increase in colocalization of LRRK2 and Rab5a in the presence of SAR405, while cells expressing LRRK2[G2019S] showed a moderate increase in colocalization with Rab5a-positive membranes (Figure 3A,C,D; Supplementary Figure 4A,B). In contrast, colocalization between LRRK2[R1441H] and Rab5a was increased upon treatment with SAR405 (Figure 3B,C,D; Supplementary Figure 4C)^24^. Co-expressing DARPin C12 with either LRRK2[WT] or LRRK2[G2019S] in the presence of SAR405 increased the colocalization between LRRK2 and Rab5a (Figure 3A,C,D; Supplementary Figure 4A). This supports our hypothesis that unlatched LRRK2 is preferentially localized to Rab5a-positive endosomes in response to SAR405. Given that LRRK2[R1441H] favors unlatching of the N-terminal repeats, we further hypothesized that the presence of DARPin C12 would not result in any additional increase in the colocalization of LRRK2[R1441H] with Rab5a-positive membranes. This was indeed the case (Figure 3A,C,D). Treating cells with SAR405 also led to an increase in Rab8a phosphorylation (Supplementary Figure 5). Our data indicates that promoting the unlatched conformation of LRRK2 increases its association with membranes.

**Figure 3.**
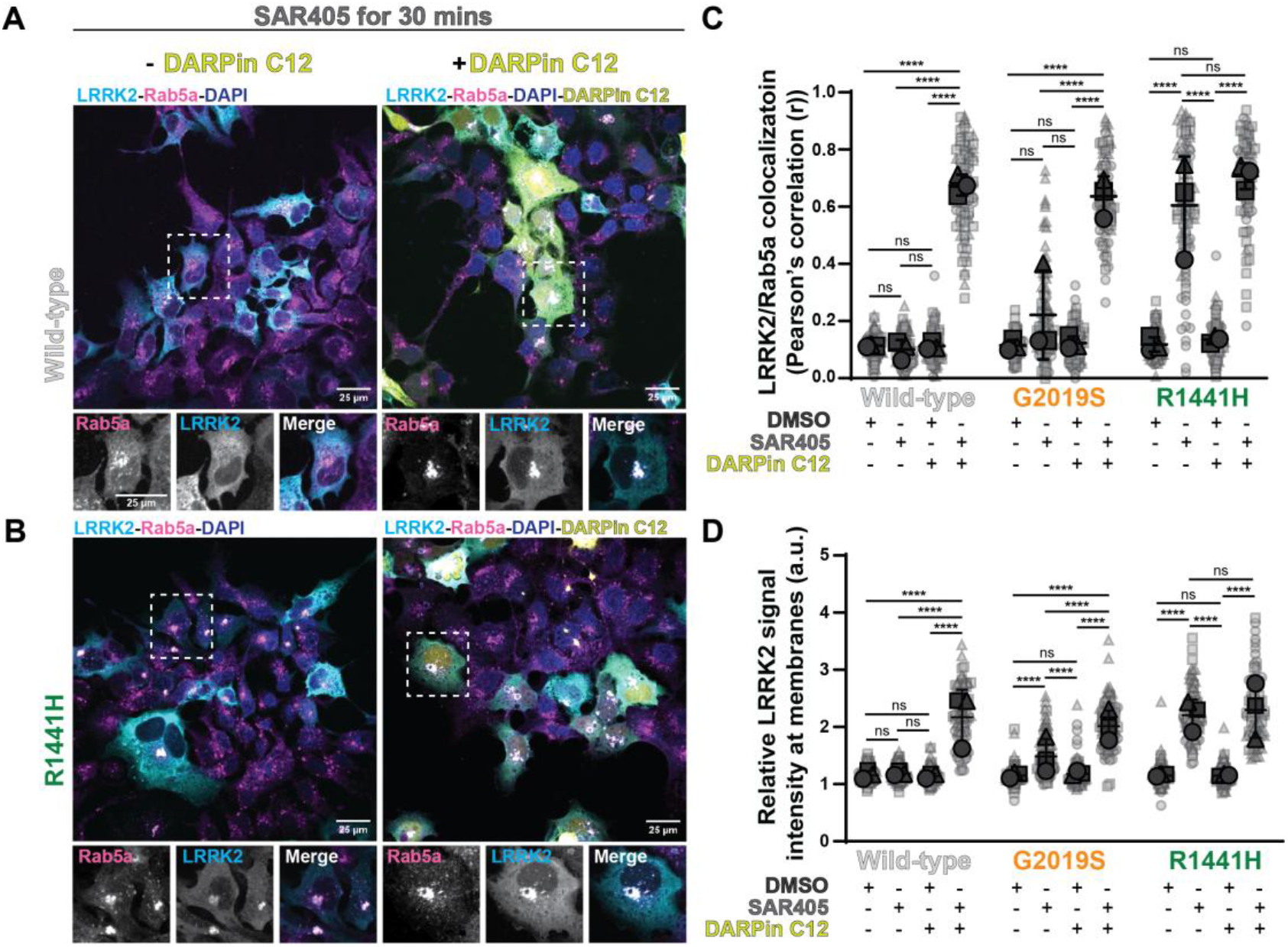
The unlatched conformation of LRRK2 is preferentially localized to membranes. **(A, B)** Representative images of 293Ts expressing GFP11-tagged LRRK2[WT] (A) or LRRK2[R1441H] (B) without (-) or with (+) GFP-tagged DARPin C12, 30 minutes after the addition of VSP34 inhibitor SAR405 (6 μM). Cells were immunostained for LRRK2, Rab5a, and GFP-DARPin C12, and nuclei visualized with DAPI. Control cells were treated with DMSO (Supplementary Figure 4B,C). Scale bars, 25 μm. **(C)** Quantification of the colocalization of GFP11-tagged LRRK2[WT] (expanded in Supplementary Figure 4B), LRRK2[G2019S] (Supplementary Figure 4A) and LRRK2[R1441H] (expanded in Supplementary Figure 4C) and endogenous Rab5a using Pearson’s correlation coefficient when cells were treated with either DMSO or SAR405. Ten images were collected per condition for each of 3 independent experiments (n=3). Three cells from each image were processed using ImageJ, and data were analyzed using a two-way ANOVA corrected for multiple comparisons using Tukey’s test. **** p≤0.0001; ns=non - significant. Error bars +/- SD. **(D)** Quantification of the ratio of mean signal intensity of LRRK2[WT], LRRK2[G2019S] and LRRK2[R1441H] at endogenous Rab5a puncta over the mean LRRK2 signal intensity of the whole cell (ROI) when cells were treated with either DMSO or SAR405. Ten images were collected per condition for each of 3 independent experiments (n=3). Rab5a positive puncta from three cells per image were identified and segmented using Ilastik. Mean signal intensity was measured using ImageJ, and data were analyzed using a two-way ANOVA corrected for multiple comparisons using Tukey’s test. **** p≤0.0001; ns=non -significant. Error bars +/- SD. Images analyzed using this approach are the same as those analyzed in Figure 3C.

### Rab8a stimulates GTP hydrolysis in unlatched LRRK2

Previously, we showed that Rab8a promotes LRRK2 GTPase activity in the presence of DARPin C12^23^. Here, we further examined the effects of repeat unlatching on LRRK2 GTPase activity *in vitro*. We measured the GTPase activity of LRRK2[WT], LRRK2[G2019S] and LRRK2[R1441H] in the presence and absence of DARPin C12 and GTPase deficient Rab8a[Q67L] (Figure 4A). For both LRRK2[WT] and LRRK2[G2019S], a combination of both C12 and Rab8a was required to activate LRRK2 GTPase activity (Figure 4B). This is consistent with our previous work on LRRK2[WT] and indicates that the LRRK2[G2019S] mutation does not impact GTPase activity. Similarly, LRRK2[G2019S] did not impact the rate of GDP release; as previously described for LRRK2[WT], addition of DARPin C12 is required to promote release of GDP from LRRK2[G2019S] (Supplementary Figure 6)^23^. However, Rab8a[Q67L] alone was sufficient to stimulate GTPase activity of LRRK2[R1441H], with no additional increase in the rate of GTP hydrolysis observed in the presence of DARPin C12 (Figure 4B). This indicates that the unlatched conformation is required for Rab8a to stimulate GTPase activity.

**Figure 4.**
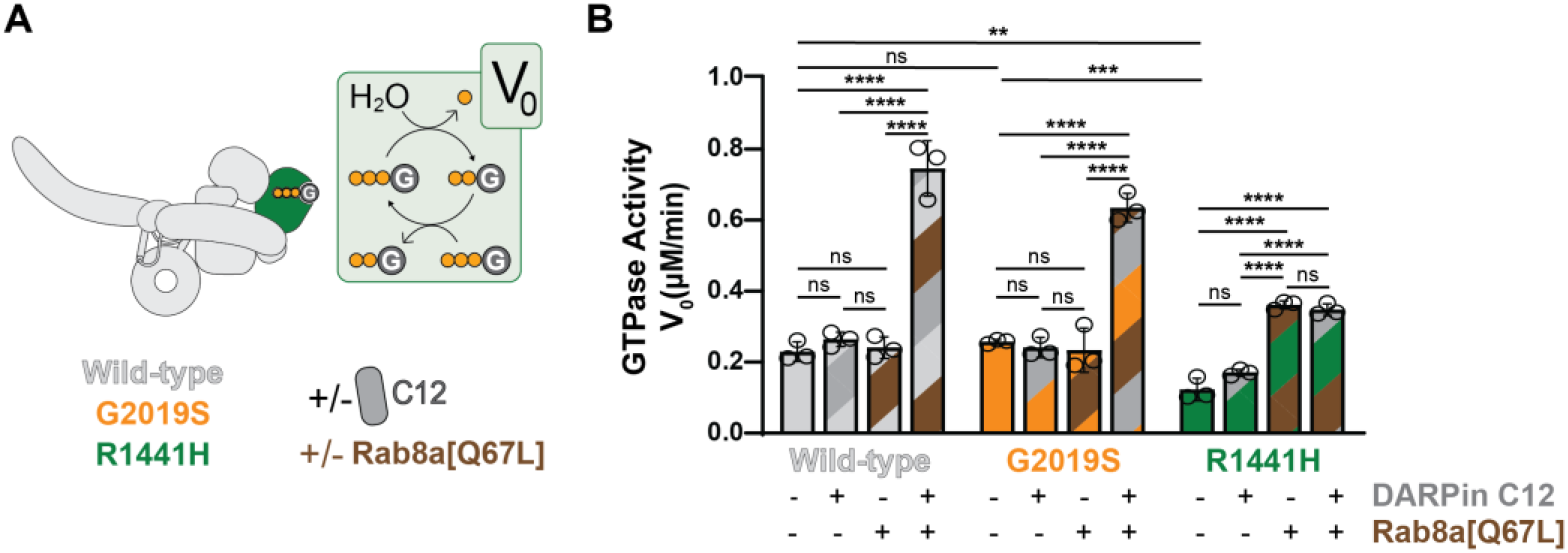
Rab8a[Q67L] stimulates GTP hydrolysis when the leucine-rich repeats are unlatched. **(A)** Schematic of the LRRK2 GTPase assay setup. LRRK2 hydrolyzes GTP via its ROC GTPase domain (green), releasing inorganic phosphate (Pi). The rate of Pi release is monitored to determine the initial reaction velocity (V_0_), which is used as a measure of GTPase activity^23^. Assays were performed with LRRK2[WT], LRRK2[G2019S], and LRRK2[R1441H] in the presence (+) or absence (-) of DARPin C12 and constitutively GTP-bound Rab8a[Q67L], as indicated. **(B)** Initial GTP hydrolysis rates (V_0_, μM/min) for LRRK2[WT], LRRK2[G2019S], and LRRK2[R1441H] under four conditions: LRRK2 alone, plus (+) DARPin C12, plus (+) Rab8a[Q67L], or plus (+) both DARPin C12 and Rab8a[Q67L]. Statistical significance was determined by a two-way ANOVA with Tukey’s multiple comparisons test: ** p≤0.01; *** p≤0.001; ****p≤0.0001; ns=non-significant. Error bars +/- SD.

As described in previous studies, LRRK2[R1441H] exhibited reduced intrinsic GTPase activity compared to both LRRK2[WT] and LRRK2[G2019S] (Figure 4B)^33,58^. Unlatching the leucine-rich repeats with DARPin C12 did not lead to reduced GTPase activity in any of the contexts that we tested, suggesting that repeat unlatching is not mechanistically responsible for the reduction in GTPase activity displayed by R1441H. The molecular basis of this remains to be determined.

### LRRK2[I2020T] is an unlatching variant

The importance of functional readouts for determining mechanism is best exemplified by the PD variant I2020T. G2019 and I2020 are consecutive residues in the kinase domain, and PDassociated mutations at these residues have highly similar structures in both the ‘activelike’ and inactive conformations (Supplementary Figure 7)^50^. Unlike LRRK2[G2019S], in cellular assays LRRK2[I2020T] can phenocopy the ‘unlatching mutants’, indicating potential similarities in mechanism^7,24,59,60^. LRRK2[I2020T] has been shown to increase both association of LRRK2 with microtubules^11,32^ and colocalization with Rab5-positive membranes under SAR405 treatment^24^. These effects mirror what is also described for mutations at R1441^10,31,23^. To determine if LRRK2[I2020T] is an unlatching variant, we tested the impact of DARPin C12 on phosphorylation of both substrate Rab8a and the newly defined biomarker for unlatching, S935. In these conditions, LRRK2[I2020T] increases phosphorylation of Rab8a compared to LRRK2[WT] but shows no further increase in the presence of DARPin C12 (Figure 5A; Supplementary Figure 8), indicative of increased substrate phosphorylation as a result of preferential repeat unlatching. Similarly to LRRK2[R1441H], DMSO treated LRRK2[I2020T] has reduced phosphorylation at S935 compared to LRRK2[WT] and LRRK2[G2019S]. Phosphorylation at this site is restored by treatment with the type II inhibitor RN277 (Figure 5B). Based on our findings that reduced phosphorylation at S935 is indicative of the unlatched state (Figure 2B), this implies that LRRK2[I2020T] favors unlatching of the leucine-rich repeats. LRRK2[I2020T] autophosphorylation however is distinct from both LRRK2[WT] and LRRK2[R1441H]. In our assay, LRRK2[I2020T] does not display increased autophosphorylation at S1292 as described for other PD-associated gain-of-function variants. However, phosphorylation at S1292 is not increased by the presence of DARPin C12 in LRRK2[I2020T], unlike LRRK2[WT] (Supplementary Figure 9). This indicates that while LRRK2[I2020T] is an unlatching mutant, the exact mechanism likely differs from the ROC:COR-B interface mutations R1441H and Y1699C.

**Figure 5.**
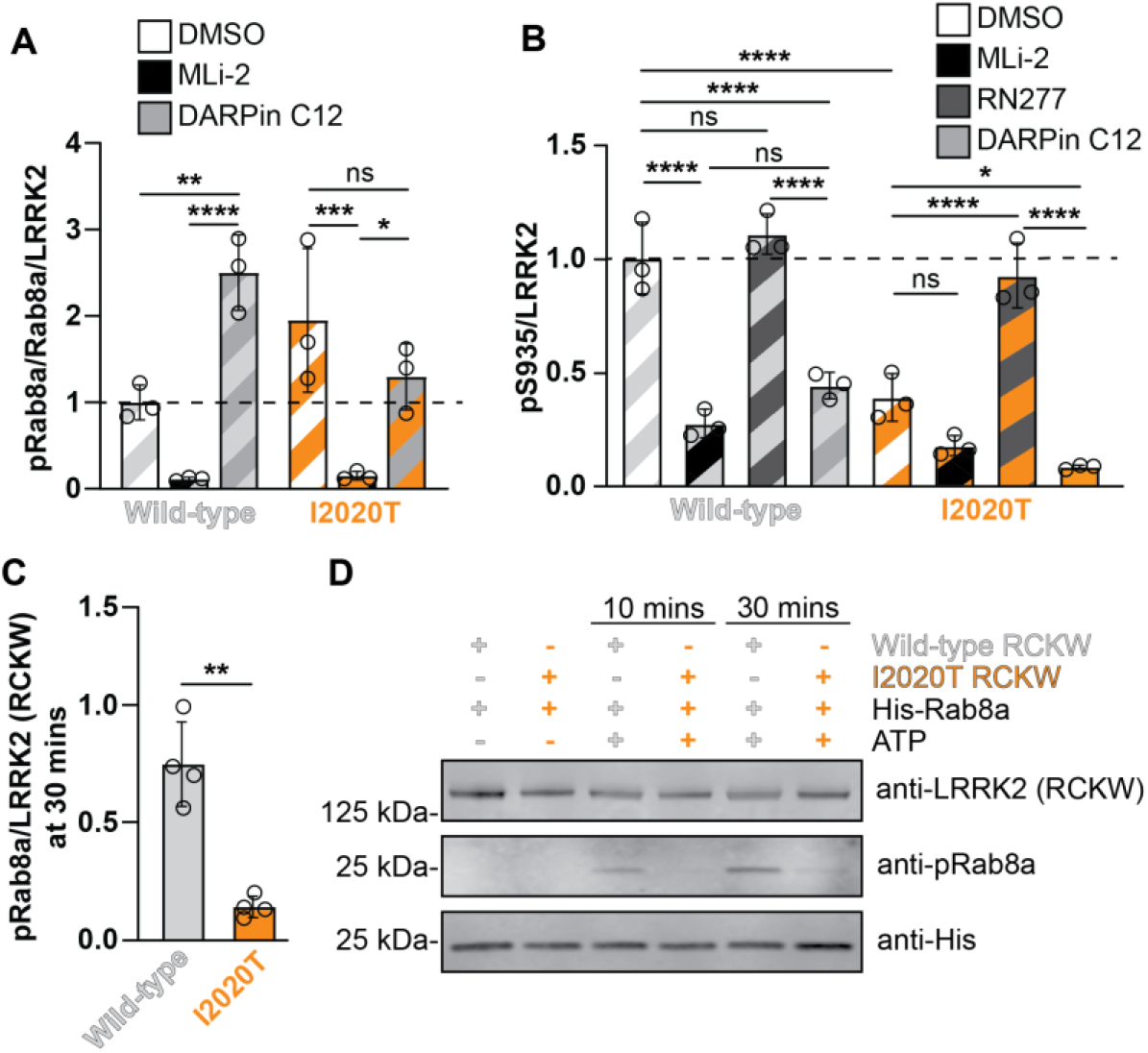
Kinase mutation I2020T is an unlatching variant. **(A)** Quantification of Rab8a (pT72)/total Rab8a/total LRRK2, normalized to the LRRK2[WT] DMSO condition from 3 (n=3) independent western blots. 293T cells overexpressing GFP11-tagged LRRK2[WT] or LRRK2[I2020T] mutant with or without FLAG-tagged DARPin C12 were treated with either DMSO or MLi-2 (5 μM, 1 hour). Blots were processed on ImageJ and data were analyzed using a two-way ANOVA with Tukey’s multiple comparison correction. * p≤0.05; ** 0.001<p≤0.01; *** p≤0.001; ****p≤0.0001; ns=non-significant. Error bars +/- SD. Representative immunoblots related to this analysis can be found in Supplementary Figure 8. **(B)** Quantification of LRRK2 (pS935)/total LRRK2 from 293T cells overexpressing GFP-11 tagged LRRK2[WT] or LRRK2[I2020T], normalized to the LRRK2[WT] DMSO condition from 3 (n=3) independent western blots. Cells were treated with DMSO, Mli-2 (1 μM), or RN277 (10 μM) for 1 hour. Blots were analyzed as previously stated. Data were analyzed using a one-way ANOVA with Tukey’s correction for multiple comparisons. * p≤0.05; *** p≤0.001; ****p≤0.0001; ns=non-significant. Error bars +/- SD. Representative immunoblots related to this analysis can be found in Supplementary Figure 9. **(C)** Quantification of the phosphorylation of recombinant His-tagged Rab8a (pT72)/RCKW, 30 minutes after the addition of ATP for four (n=4) independent *in vitro* kinase assays. Purified RCKW[WT] or RCKW[I2020T] (100 nM) was incubated with His-tagged Rab8a (1 μM) and ATP (2 μM) for either 10 or 30 minutes. No ATP was used as a control. Samples were immunoblotted for LRRK2, His, and Rab8a (pT72) and processed on ImageJ. Data were analyzed using Welch’s t-test. ** 0.001<p≤0.01. Error bars +/- SD. **(D)** Representative immunoblot for Figure 5C.

Given the position of I2020T in the kinase active site, adjacent to the activation loop, we hypothesized that the mutation may confer additional effects beyond favoring conformational unlatching. To test for an effect on kinase activity independent from repeat unlatching, we performed *in vitro* kinase activity assays in the absence of the N-terminal repeat domains (residues 1227-2527: ‘RCKW’). We found that RCKW[I2020T] phosphorylated substrate Rab8a to a lesser extent than RCKW[WT] (Figure 5C,D). Reduced kinase activity of RCKW[I2020T] compared to RCKW[WT] is consistent with previous reports^61^. We therefore define I2020T as a dual-effect mutation, promoting the unlatched conformation to increase substrate phosphorylation in the presence of the N-terminal repeats, but intrinsically reducing the activity of the kinase.

## Discussion

Maintaining the autoinhibited state of LRRK2 is a critical intramolecular mechanism for regulating activity, and a key point of pathogenic dysregulation in PD. In this study, we identify unlatching of the leucine-rich repeats as a shared pathogenic mechanism leading to increased substrate phosphorylation across multiple PD-associated variants. Notably, G2019S is the only mutation that we tested that does not stimulate repeat unlatching. The ROC:COR-B interface is central to stabilizing LRRK2’s autoinhibited conformation, as structurally characterized in our previous work^23^, and its role in maintaining autoinhibition provides a mechanistic explanation for the hotspot of PD-associated mutations in this region. Our data also provides a framework for stratifying the mechanism of action of PD-associated mutations that cannot be readily inferred from structural analysis, as we demonstrate for LRRK2[I2020T]. Despite its proximity to the G2019S mutation in the kinase and no structural differences from LRRK2[WT] or LRRK2[G2019S] (Supplementary Figure 7)^50^, our cellular assays indicate that LRRK2[I2020T] promotes unlatching of the leucine-rich repeats. This mechanistically explains the phenotypic similarities between I2020T and our newly defined category of ‘unlatching mutations’ described across multiple previous studies from our lab and others^7,11,24^. Furthermore, we show that the activating effect of the I2020T mutation is dependent on the presence of the N-terminal repeat domains, consistent with previous reports^61^. We propose that I2020T is a dual-effect mutation, facilitating repeat unlatching while directly impairing intrinsic catalytic activity. The balance of these opposing effects may explain the conflicting reports in the literature relating to the impact of I2020T on kinase activity, based on experimental set up, construct design and readout^7,59,62^. By defining the molecular and cellular consequences of unlatching the leucine-rich repeats, we expand the current model of LRRK2 activation and increase mechanistic understanding of how activation/autoinhibition are dysregulated in disease (Figure 6). Given the variability in penetrance and clinical manifestations of different PD-associated mutations, understanding how individual variants perturb the regulatory mechanisms of LRRK2 may be important in the development of diagnostic tests and novel, mutation-specific therapies.

**Figure 6.**
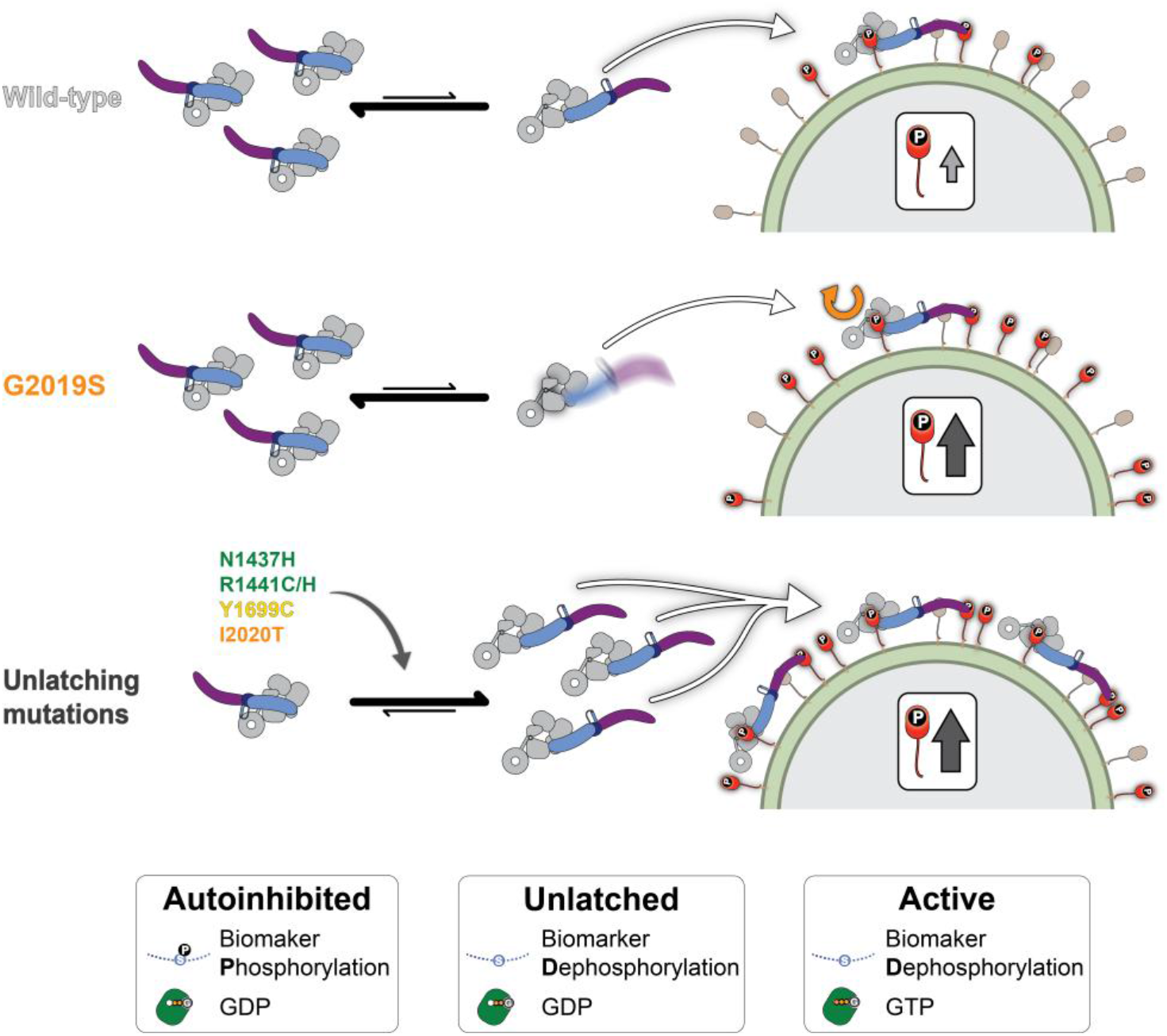
Proposed model of LRRK2 activation and dysregulation by PD-associated mutations. The majority of the cytosolic population of LRRK2[WT] and [G2019S] is in the autoinhibited conformation, with the leucine-rich repeats latched across an open, inactive kinase. In this conformation, the phosphorylation site S935 on the disordered loop preceding the LRR domain is protected from dephosphorylation, likely due to association with 14-3-3 family proteins^27^, and the ROC domain is bound to GDP. For LRRK2 to be activated and phosphorylate substrate, the leucine-rich repeats must unlatch from their position occluding the kinase active site. In the unlatched state, the steric constraints maintaining the kinase in its open, inactive conformation are relieved, the affinity for GDP in the ROC domain is reduced, and S935 is exposed for dephosphorylation. ‘Unlatching mutations’ perturb the conformational equilibrium between the autoinhibited and unlatched states, favoring repeat unlatching. In contrast, LRRK2[G2019S] does not affect this step of the activation cycle. Spatially, LRRK2 phosphorylates its substrate Rab GTPases on intracellular membranes. Unlatching mutants increase localization of LRRK2 on membranes. Here, stimulated by the presence of Rab substrate, nucleotide exchange from GDP to GTP stabilizes the active conformation; this stably active conformation is responsible for the majority of Rab substrate phosphorylation ^23^. Mutations such as G2019S that likely increase the inherent catalytic rate of the kinase act on this step.

### Phosphorylation at S935 is a biomarker for the unlatched conformation, not kinase activity

Destabilizing the ROC:COR-B interface promotes unlatching of the leucine-rich repeats, priming LRRK2 for substrate phosphorylation^12,23^. Conversely, stabilizing the ROC:COR-B interface and/or the latched conformation of the leucine-rich repeats would reduce LRRK2 activity by sterically constraining the kinase in the inactive state. Previous studies indicate that this is the mechanism employed by the best-established regulators of LRRK2, the 14-3-3 family proteins. As demonstrated in a recently published structure of the 14-3-3:LRRK2 complex, dimeric 14-3-3 interacts with LRRK2 in two places: a pair of phosphorylated serine residues (S910 and S935) within a flexible loop preceding the LRR domain, and the COR subdomains^27,52,63^. Interaction of 14-3-3 with the latter region is only possible when LRRK2 is in the autoinhibited conformation^27^. Dephosphorylation of the 14-3-3 recruitment sites, specifically S935, has been widely interpreted as a proxy for kinase activity, and used to assess the efficacy of LRRK2 inhibitors in cells, animal models and clinical trials^48,49,64,65^. However, our data, supported by contributions from several other groups^7,27,30^, clarifies that phosphorylation at these sites should be interpreted as a readout of conformation rather than kinase activity. Dephosphorylation at S935 correlates with unlatching of the leucine-rich repeats, induced by both ‘unlatching mutations’ and DARPin C12 that increase LRRK2 substrate phosphorylation, and type I kinase inhibitors, which prevent it. Conversely, stabilizing the autoinhibited conformation with type II kinase inhibitors restores phosphorylation at this site.

Future work should clarify, mechanistically, how phosphorylation of S935 is regulated in a cellular context. Does the conformation-specific association of 14-3-3 to LRRK2 in the autoinhibited state determine phosphosite accessibility to upstream phosphatases such as PP1 and PP2A? Specifically, does 14-3-3 binding shield these phosphosites in the autoinhibited state and expose them when 14-3-3 dissociates from the unlatched conformation?

This hypothesis is supported by several previous studies which show that phosphorylation of S935 is reduced when 14-3-3 binding to LRRK2 is prevented^63,66^. Alternatively, are kinases and phosphatases recruited to LRRK2 in a conformation-specific manner, or does conformation of LRRK2 itself regulate access of upstream regulators to these phosphorylation sites? The loop containing S935 is highly flexible and therefore disordered in all published structures of LRRK2 to date^12,23,27,47^; this hypothesis therefore seems the least likely, although it is possible that a key conformation of LRRK2 is yet to be observed in structural studies.

Interpretation of S935 phosphorylation as a biomarker of conformational state has potential to provide insight into disease mechanisms of a broad group of PD patients beyond those carrying a mutation in LRRK2. Increased phosphorylation of LRRK2 substrates has been observed in idiopathic cases of PD^8–10^ and carriers of non-LRRK2 mutations, most clearly VPS35 D620N^67^. This suggests a role for LRRK2, although the mechanism behind increased LRRK2 activity in these contexts is unknown. Stratifying PD patients with elevated Rab phosphorylation by pS935 level could distinguish LRRK2 dysregulation driven by unlatching from that driven by other, as yet uncharacterized, mechanisms.

### Unlatched LRRK2 preferentially localizes to membranes

In our model, LRRK2 associates with membranes in the unlatched state. Membrane recruitment is a key step in the cellular activation of LRRK2, bringing it into proximity with its membrane-bound Rab GTPase substrates^54,61,68^. LRRK2 recruitment to membranes can be induced through multiple cellular pathways in response to a variety of upstream stimuli including lysomotropic drugs^55^, cytoplasmic DNA^56^, and endosome/phagosome membrane damage by intracellular pathogens^57^. We show that LRRK2 localizes to Rab5a-positive vesicles upon treatment with the VPS34 inhibitor SAR405, and our data suggest that unlatching of the leucine-rich repeats promotes this membrane association. However, our work cannot distinguish whether unlatching affects conformation-specific recruitment or dissociation. LRRK2 is recruited to membranes by recruitment factors^35,69^, the best characterized of which are non-substrate Rab GTPases. Three Rab-binding sites in the LRRK2 ARM domain have been identified and functionally implicated in recruiting LRRK2 to organelle membranes^35^. Whether recruitment is sensitive to conformational state remains to be determined, but structural studies^35^ show that these known recruitment sites are likely equally accessible in both the autoinhibited and unlatched states. There is therefore no apparent physical reason why recruitment factors would favour the unlatched conformation. Alternatively, autoinhibited LRRK2 could be preferentially maintained in the cytosol by additional factors, although other than 14-3-3, none have yet been identified.

Increased membrane localization of unlatched LRRK2 could also result from stabilization of the unlatched, membrane-bound pool. Phosphorylated substrate Rab GTPases are likely to contribute to this. Previous work shows that reducing the pool of phosphorylated Rab GTPases reduces the amount of LRRK2[R1441C] localized on Rab5a-positive vesicles following SAR405 treatment^24^. This indicates that association of LRRK2 with membranes requires phosphorylated Rab GTPases, as proposed in the feed-forward model of LRRK2 membrane recruitment^69^. Destabilizing the autoinhibited state, through gain-offunction mutations or DARPin C12, also increases Rab substrate phosphorylation. It is likely that this increase in the pool of phosphorylated Rabs contributes to improved membrane association, and this may explain the slight increase in colocalization of LRRK2[G2019S] and Rab5a upon SAR405 treatment compared to LRRK2[WT] in our study. Finally, unlatched LRRK2 may make additional interactions with the membrane or membrane-bound cofactors, though no such conformation-specific interactions have yet been identified.

## Data availability

All data needed to evaluate the conclusions of the study are present in the paper and/or the Supplementary Materials. All other data supporting the findings of this study are available from the corresponding authors upon reasonable request. All reagents used in this study will be shared upon request.

## Acknowledgements

The authors would like to thank all members of the Reck-Peterson and Leschziner labs for their feedback and advice during the preparation of this manuscript. The authors would also like to thank their funding sources for supporting this work (listed below).

## Author contributions

This study was conceptualized by K.J.S. and S.L.R-P. K.S.H., K.J.S., and W.G. designed the experiments. K.S.H., K.J.S., and B.Q. performed the cell-based activity assays. W.G. performed the GTPase and GDP release assays. K.S.H., and B.Q. performed the imaging experiments, supported by G.K. and N.L. R.A. and N.P. provided the GFP11-LRRK2 construct. K.S.H., K.J.S. and W.G. analyzed the data. K.S.H. and K.J.S. wrote the original manuscript, and S.L.RP reviewed and edited it.

## Materials and Methods

### Plasmid generation and cloning for mammalian cell expression

All LRRK2 mutant constructs for mammalian cell expression were generated by Genscript. The pcDNA5/FRT TO GFP11-LRRK2 plasmid (RRID: Addgene_231174) was used as a template for site-directed mutagenesis. All plasmids used in this study were validated by whole-plasmid sequencing.

### Mammalian cell culture

All cellular assays were conducted using 293T HEKs (ATCC, RRID: CVCL_0063). Cells were cultured in Dulbecco’s Modified Eagle Medium (Corning, DMEM) supplemented with 10% Fetal Bovine Serum and 1% Penicillin-streptomycin (Corning) at 37°C in 5% CO2. For western blot and imaging experiments respectively, cells were either plated into tissue culture treated 6-well plates (CELLTREAT) at a concentration of 0.3×106 cells per well, or onto fibronectin-coated (Sigma-Aldrich, catalog #F0895) glass coverslips at a concentration of 0.2×106 cells per well and incubated at 37°C in 5% CO2. All compounds used to treat mammalian cells were prepared in DMSO.

### Transfection of mammalian cells

Cells were incubated at 37°C in 5% CO_2_ for 48 hours after plating, then transfected with the indicated constructs using polyethylenimine (Polysciences, catalog #23966-1) at a 3:1 PEI:DNA ratio unless otherwise described.

For analysis of Rab8a phosphorylation, cells were transfected with 1000 ng of GFP11-tagged LRRK2[WT], [R1441H], [G2019S], [Y1699C], [N1437H], [L1795F], or [I2020T] and 500 ng GFP-Rab8a (Addgene, RRID: Addgene_49543), with or without 500 ng of 8xHis-DARPin C12-3xFLAG (pcDNA3.1 vector backbone)^23^. After incubating for a further 48 hours, cells were treated with either MLi-2 (Cayman Chemical Company, catalog #19305; 5 μM) or DMSO and incubated for one hour at 37°C in 5% CO_2_ before lysis.

For analysis of LRRK2 phosphorylation, cells were transfected with 800 ng of GFP11-LRRK2[WT], [R1441H], [G2019S] or [I2020T] with or without 400 ng of DARPin C12. After incubating for 48 hours, cells were treated with either MLi-2 (1 μM), RN277^47^ (10 μM), or DMSO and incubated for three hours at 37°C in 5% CO2 before lysis.

For analysis of LRRK2 activation after SAR405 treatment, cells were transfected with 1000 ng of GFP11-tagged LRRK2[WT] or LRRK2[R1441H]. After 48 hours, cells were treated with MLi-2 (1 μM, 1 hour), SAR405 (MedChemExpress, catalog #HY-12481; 6 μM, 30 minutes), or DMSO and incubated at 37°C in 5% CO_2_ before lysis.

For immunofluorescence imaging experiments, cells were transfected 24 hours after plating with 800 ng of GFP11-LRRK2[WT], [R1441H] or [G2019S] with or without 400 ng of DARPin C12 using FuGENE HD Transfection Reagent (Promega, catalog #E2311). After a further 24 hours, cells were treated with either DMSO or SAR405 (6 μM) for 30 minutes at 37°C in 5% CO_2_.

### Immunoblotting and analysis for cellular experiments

Transfected 293T cells were washed three times in cold DPBS before lysis of each well in 300 µl RIPA buffer (50 mM Tris pH 8, 150 mM NaCl, 1% Triton X-100, 0.1% SDS, 0.5% Sodium Deoxycholate, 1 mM dithiothreitol (DTT)) in the presence of cOmplete mini EASYpack protease inhibitor cocktail (Sigma-Aldrich, catalog #05056489001) and PhosSTOP EASYpack phosphatase inhibitor cocktail (Roche, catalog #04906837001). Lysates were then vortexed for 30 seconds and slowly agitated for 15 minutes at 4°C. After removing cellular debris by centrifugation at 17,000 x g for 15 minutes at 4°C, supernatant was moved to a new tube and gel samples were prepared by adding SDS-PAGE loading buffer and reducing agent (1X, Invitrogen, catalog #NP0007 and NP0009). Samples were boiled at 95°C for 10 minutes and then either stored at −20 °C or run immediately on a 4-12% Bis-Tris gel (Invitrogen). Samples were immunoblotted as described previously^37^. Briefly, proteins were transferred onto PVDF membrane (Thermo Scientific, catalog #88520) at 200 mA for 4 hours at 4 °C. Following transfer and blocking in 5% milk or BSA for 1 hour at room temperature, membranes were cut above the 70 kDa marker (Chameleon Duo prestained protein ladder; LiCORbio, catalog #928-60000). The top portion of the membrane was immunoblotted either for LRRK2 alone with a LRRK2 specific antibody (MJFF2-c41-2, RRID: AB_2713963) in 5% BSA or LRRK2 (N241A/34, RRID: AB_2877351) multiplexed with either phospho-LRRK2 S935 (UDD2 10(12), RRID: AB_2904231) or phospho-LRRK2 S1292 (MJFR-19-7-8, RRID: AB_2921223) in 5% milk. The lower portion of the membrane was immunoblotted with antibodies for GFP (RRID: AB_627695), phospho-Rab8a T72 (MJF-R20, RRID: AB_2814988), and GAPDH (RRID:AB_561053) in 5% milk, or cut again at around the 30 kDa marker and probed for GAPDH (upper) in 5% milk and Rab10 (605B11, RRID: AB_2921226) and phospho-Rab10 T73 (lower) (MJF-R21, RRID: AB_2811274) in 5% BSA. Antibodies raised in different species were multiplexed as indicated in the text and figure legends. For all primary antibodies, membranes were incubated with the antibody dilutions overnight at 4 °C before washing three times for 10 mins with Tris-buffered saline supplemented with 0.1% Tween-20 (TBS-T). Washed blots were subsequently incubated for 1 hour at room temperature with secondary antibodies (IRDye 680RD Goat Anti-Rabbit, RRID: AB_2721181 and IRDye 800CW Donkey Anti-Mouse, RRID: AB_2716622). Blots were washed a further three times in TBS-T and imaged. Following visualization, membranes were either subsequently cut at 25 kDa marker and immunoblotted for FLAG with a FLAG-specific antibody (PGT Labs, RRID:AB_11232216), or samples were loaded identically and run on Tricine 10%-20% gels (Invitrogen) as described previously, to visualize FLAG-DARPin C12 by immunoblotting with an anti-FLAG antibody^23^. Downstream transfer and immunoblotting of Tricine gel samples was performed as for Bis-Tris gels described above. Antibodies and working concentrations are listed in Supplementary Table 1. All membranes were imaged using the LI-COR imaging system. Intensities of probed protein bands were measured using ImageJ (version 2.14.0). Statistical tests were performed as indicated in the text and figure legends using GraphPad Prism (version 10.5.0, RRID: SCR_002798).

### Immunofluorescence, confocal microscopy and analysis

Cells were washed on ice with cold DPBS three times, then incubated at room temperature for 10 minutes in 4% PFA/3% sucrose fixation buffer. Fixed cells were washed another four times with cold DPBS, before being incubated with a combined permeabilization, blocking and staining buffer (5% Donkey Serum, 0.2% Saponin, 0.1 M PHEM buffer (Electron Microscopy Sciences, catalog #11165) in DPBS) with primary antibodies against Rab5a (Invitrogen, RRID:AB_2884043) and LRRK2 (Abcam UDD3 30(12), RRID:AB_2938806) for 1 hour at room temperature. All antibodies for immunofluorescence can be found with details of working concentrations in Supplementary Table 1. Cells were then gently washed four times with cold DPBS before incubation with secondary antibodies (Alexa Fluor 586 Goat Anti-Mouse IgG (H+L), Alexa Fluor 647 Chicken Anti-Rabbit IgG (H+L), respectively) and DAPI (Sigma Aldrich, catalog #D9542; 300 nM) for 30 minutes at room temperature in the dark. Cells were washed three times in DPBS before being mounted on coverslips with FluorSave reagent (EMD Millipore, catalog #345789) and left to cure in the dark for 24 hours. Samples were then blinded and imaged on a Yokogawa W1 confocal scan head mounted to a Nikon Ti2 microscope with an Apo ×60 1.49-NA objective. The microscope was run with NIS Elements using the 488 nm, 561 nm, 640 nm and 405 nm lines of a six-line LUN-F-XL laser engine and a Prime95B camera (Photometrics). Ten images were taken per coverslip. Images were first analyzed blind for colocalization using Pearson’s correlation using ImageJ (version 2.14.0). Three ROIs were selected per image by tracing the outline of an individual cell using saturated LRRK2 signal as a guide to define cell boundaries. As the cells were overexpressing LRRK2, the background was then subtracted from each ROI using a rolling ball radius of between 5 and 10, and the outside of the ROI was cleared. ROIs were then thresholded using the default setting reduced to between 1-10%. Colocalization between the resulting masks of the LRRK2 and Rab5a channels were reported using Pearson’s correlation coefficient from the JACoP (version 2.1.1) colocalization plugin. Additionally, ROIs from the same images were analyzed for the ratio of mean LRRK2 signal intensity at Rab5a puncta to the mean LRRK2 signal intensity of the whole ROI. Three to five ROIs per condition were used to manually train Ilastik’s machine learning segmentation software (version 1.4.2) on the color/intensity, edge, and texture (Sigma=3.5) of Rab5a positive puncta. Simple segmentation of the Rab5 puncta in each ROI were downloaded and converted to masks on ImageJ. Mean LRRK2 signal intensity at the Rab5a puncta and across the whole ROI were measured. All statistical tests were applied as indicated in the text and performed using GraphPad Prism (version 10.5.0, RRID: SCR_002798). Samples remained blinded throughout both imaging and analysis.

### Expression and purification of LRRK2 and RCKW[WT] from insect cell expression system

For LRRK2 expression, DNA encoding full-length LRRK2[WT] (residues 1– 2527) or RCKW LRRK2 (residues 1327-2527) from the Mammalian Gene Collection was amplified via polymerase chain reaction (PCR) and inserted into the pKL expression vector (Addgene, RRID: Addgene_110741). The full length LRRK2 plasmid served as the template for introducing LRRK2[G2019S], [I2020T], and [R1441H] mutations using the Q5 Site-Directed Mutagenesis Kit (NEB, catalog #E0554S). The resulting constructs were used to generate recombinant baculoviruses following the Bac-to-Bac expression system protocols (Invitrogen) as previously described^23^. Sf9 insect cells (RRID:CVCL_0549) were then infected with baculovirus and cultured at 27 °C for 3 days in SF-900II SFM (Gibco) media. 3 days post-infection, cells were collected and cell pellets were resuspended in lysis buffer (50 mM HEPES pH 7.4, 500 mM NaCl, 20 mM imidazole, 0.5 mM tris(2-carboxyethyl) phosphine (TCEP), 5% glycerol, 5 mM MgCl2, 20 μM GDP, 0.5 mM protease inhibitor cocktail (Roche)). Cells were lysed using a Dounce homogenizer and the lysate clarified by centrifugation. The supernatant was removed to a new tube and incubated for 1 hour with Ni-NTA agarose beads (Qiagen) pre-equilibrated in lysis buffer. Beads were applied to a gravity column where they were extensively washed with lysis buffer, followed by elution in lysis buffer supplemented with 300 mM imidazole. The eluate was incubated at 4 °C overnight with 1 µM recombinantly expressed Tobacco Etch Virus (TEV) protease to remove the tag. The protein sample (containing cleaved protein, cleaved tag, TEV protease and other contaminating proteins) was concentrated and loaded directly onto a S200 size exclusion column (Cytiva) pre-equilibrated in storage buffer (50 mM HEPES pH 7.4, 500 mM NaCl, 0.5 mM TCEP, 5% glycerol, 5 mM MgCl2 and 20 μM GDP) to remove contaminants. Purified LRRK2 was concentrated to ∼5–6 μM and flash frozen in liquid nitrogen for storage at −80°C. Protein purity was assessed by SDS-PAGE, and concentration was determined by absorbance at 280 nm. N-terminally tagged His6–Z–TEV–LRRK2[WT], [G2019S] and [R1441H] were purified using the same protocol.

### Expression and purification of RCKW[I2020T] from insect cell expression system

Purification of RCKW[I2020T] was performed as described previously11. Briefly, N-terminally 6x-His-Z-tagged RCKW[I2020T] was expressed in Sf9 insect cells infected with baculovirus and grown for three days at 27 °C. After pelleting by centrifugation, the cells were resuspended in lysis buffer (50 mM HEPES pH 7.4, 500 mM NaCl, 20 mM imidazole, 0.5 mM tris(2-carboxyethyl) phosphine (TCEP), 5% glycerol, 5 mM MgCl2, 20 μM GDP, 0.5 mM protease inhibitor cocktail (Roche)) and lysed by Dounce homogenization. After centrifugation, the clarified lysate was incubated with Ni-NTA agarose beads (Qiagen) and washed with lysis buffer before elution with buffer containing 300 mM imidazole. RCKW[I2020T] was diluted with buffer containing no NaCl and loaded onto a SP Sepharose column (Cytiva). Protein was eluted with a 250 mM to 2.5 M NaCl gradient and then cleaved by a TEV protease overnight at 4 °C. Cleaved RCKW[I2020T] was then isolated with a second Ni-NTA column, then concentrated and run on a S200 size exclusion column equilibrated with storage buffer (20 mM HEPES pH 7.4, 700 mM NaCl, 0.5 mM TCEP, 5% glycerol, 2.5 mM MgCl2, 20 μM GDP). Protein was flash frozen and stored at -80 °C. Protein purity was assessed by SDS-PAGE, and concentration was determined by absorbance at 280 nm.

### Expression and purification of Rab8 from bacterial expression system

The coding sequence for human Rab8a residues 1-181 was cloned into a pET vector encoding a C-terminal TwinStrep and 8xHis tag and codon optimized for bacterial expression by GenScript. Plasmids were transformed into BL21 DE3 cells (NEB, catalog #C2527H). Small scale cultures were grown at 37 °C in Lysogeny broth medium (LB; Fisher Scientific, catalog #BP1426-2) supplemented with 100 µg/mL carbenicillin and used to inoculate a large-scale culture at a 1:50 dilution. At OD600 = 0.8, cultures were cooled to 18 °C and protein expression was induced with 1 mM IPTG for 16 hours at 18 °C shaking. Cells were collected into pellets by centrifugation (5,000 × g, 15 min, 4°C), media removed and the pellet flash frozen at −80 °C. On the day of purification, pellets were thawed and resuspended in 40 mL lysis buffer (50 mM HEPES pH 7.4, 300 mM NaCl, 0.5% Triton-X, 10% glycerol, 1 mM DTT, 1 mM EDTA, 20 mM imidazole, 2 mM TCEP, 2 mM PMSF) supplemented with protease inhibitor cocktail (Roche). Cells and lysis buffer were incubated rotating at 4 °C for 1 hour before being lysed by sonication. Lysate was clarified by centrifugation at 40,000 × g for 1 h at 4 °C to remove cell debris. Clarified lysate was applied to 1 mL PureCube Ni-Indigo agarose (Cube Biotech, catalog #75103) and incubated rotating at 4 °C for 1 hour before loading to a gravity flow column and washing with 25 column volumes of lysis buffer and 10 column volumes of wash buffer (50 mM HEPES pH7.4, 150 mM NaCl, 0.1% Triton-X, 10% glycerol, 50 mM imidazole, 2 mM TCEP). His tagged protein was eluted in 5 mL of elution buffer (50 mM HEPES pH7.4, 150 mM NaCl, 400 mM imidazole, 10% glycerol, 1 mM TCEP), flash-frozen in liquid nitrogen, and stored at −80 °C. Protein purity was assessed by SDS-PAGE, and concentration was determined by absorbance at 280 nm.

### Expression and purification of Rab8[Q67L] from bacterial expression system

The human Rab8a coding sequence bearing the Q67L substitution was amplified by polymerase chain reaction (PCR) and cloned into a pET-28a vector encoding an N-terminal 6×His tag and MBP tag followed by a TEV protease cleavage site (Addgene, RRID:Addgene_194702). Plasmids were transformed into *E*.*coli* Rosetta cells (Novagen, catalog #70954). Small scale cultures were grown at 37 °C in Lysogeny broth medium (LB; Fisher Scientific, catalog #BP1426-2) supplemented with 50 µg/mL kanamycin and used to inoculate a large-scale culture at a 1:1000 dilution. At OD600 = 0.6–0.8, cultures were cooled to 18 °C and protein expression was induced with 0.5 mM IPTG for 16 hours at 18 °C. Cells were collected into pellets by centrifugation (5,000 × g, 15 min, 4°C) and stored at −80 °C after the removal of media. For purification, cell pellets were resuspended in lysis buffer (50 mM HEPES pH 7.4, 500 mM NaCl, 5 mM MgCl2, 10 µM GDP, 50 mM imidazole, 0.5mM TCEP, 1 mM PMSF), incubated on ice for 30 minutes, and lysed by sonication. Clarified lysate (centrifuged at 40,000 × g, 45 min, 4 °C to remove cell debris) was applied to Ni-NTA agarose, washed with lysis buffer supplemented with 50 mM imidazole, and eluted in lysis buffer supplemented with 300 mM imidazole. The His tag was removed by TEV protease digestion (1:50 w/w) during overnight dialysis at 4 °C against 50 mM HEPES pH 7.4, 500 mM NaCl, 5 mM MgCl2, 10 µM GDP, 50 mM imidazole, 0.5mM TCEP. The dialyzed protein was further purified by size exclusion chromatography (SEC) on a Superdex 75 16/60 column (Cytiva) equilibrated in 20 mM HEPES pH 7.5, 150 mM NaCl, 5 mM MgCl2, and 1 mM DTT. Fractions were pooled, concentrated to 20 mg/mL, flash-frozen in liquid nitrogen, and stored at −80 °C. Protein purity was assessed by SDS-PAGE, and concentration was determined by absorbance at 280 nm.

### Expression and purification of DARPin C12 from bacterial expression system

DARPin C12 was expressed and purified from a pQE30 vector (Qiagen) containing an N-terminal 8× His tag and a C-terminal 3x-FLAG tag, as previously described^23^. To summarize, plasmids were transformed into *E*.*coli* Rosetta cells and grown at 37 °C with shaking until OD_600_ reached 1.0. The temperature was then reduced to 18 °C and protein expression was induced by adding 0.5 mM IPTG. After 18 hours the cells were harvested by centrifugation and resuspended in lysis buffer (50 mM HEPES pH 7.4, 500 mM NaCl, 20 mM imidazole, 0.5 mM TCEP, and 5% glycerol) prior to lysis by sonication. The lysate was clarified by centrifugation and loaded onto Ni-NTA Sepharose beads (GE HealthCare) pre-equilibrated in lysis buffer. The beads were washed with 50 column volumes of lysis buffer and protein was eluted in lysis buffer supplemented with 300 mM imidazole. The eluate was concentrated to 5 ml volume and subjected to SEC using an S190 16/200 column (GE HealthCare) in SEC buffer (20 mM HEPES pH 7.4, 150 mM NaCl, 0.5 mM TCEP, and 5% glycerol). Fractions were pooled, flash-frozen in liquid nitrogen, and stored at −80 °C. Protein purity was assessed by SDS-PAGE, and concentration was determined by absorbance at 280 nm.

### In vitro GTPase activity assay

GTP hydrolysis was monitored in real time using a colorimetric inorganic phosphate detection assay (EnzCheck Phosphate Assay Kit, Thermo Fisher Scientific/Invitrogen, E12056) as previously described^23^. In this coupled assay, released inorganic phosphate drives the purine nucleoside phosphorylase (PNP)-dependent conversion of 2-amino-6-mercapto-7-methylpurine riboside (MESG) to 2-amino-6-mercapto-7-methylpurine and ribose-1-phosphate, yielding a continuous spectrophotometric signal at 360 nm. For DARPin-only assays, purified LRRK2 (WT, G2019S, I2020T, or R1441H; 1 µM final) was incubated with or without C12 (1.2 µM final) for 30 minutes at room temperature prior to initiating GTP hydrolysis measurements. For Rab8a-dependent assays, LRRK2 (1 µM) was pre-incubated with C12 (1.2 µM) for 30 minutes at room temperature, followed by addition of Rab8a[Q67L] (20 µM) and a further 10 minutes incubation at room temperature before measurement. Absorbance at 360 nm was recorded every 5 seconds for 2 minutes using a Cary 60 UV-Vis spectrophotometer (Agilent).

### In vitro guanine nucleotide exchange assay

Nucleotide exchange of LRRK2 was measured using a fluorescence-based assay as previously described^23,58^. Recombinant LRRK2 proteins (WT, G2019S, I2020T, and R1441H; 1 µM final concentration) were preloaded with 1 mM BODIPY-FL–GDP (Thermo Fisher Scientific, catalog #G22360) in an exchange buffer consisting of 20 mM HEPES pH 7.2, 150 mM KCl, 5% glycerol, and 1 mM DTT. This loading reaction was incubated at 4 °C for 5 hours and subsequently terminated by adding MgCl_2_ to a final concentration of 10 mM. To initiate nucleotide exchange, the BODIPY-FL– GDP–loaded LRRK2 was rapidly mixed with DARPin C12 (1.2 µM) and/or Rab8a[Q67L] (20 µM) in exchange buffer supplemented with 5 mM GTP. The reactions were performed in black 384-well plates, and fluorescence intensity was monitored in real time using a microplate reader (Cytation 5, Agilent BioTek) with excitation and emission wavelengths of 488nm and 535nm, respectively. Data were recorded every 4 seconds for a total duration of 96 seconds.

### In vitro kinase activity assay with RCKW

*In vitro* kinase activity of RCKW was assessed as described in previously^61^. Briefly, purified RCKW[WT] or RCKW[I2020T] (100 nM) was combined with Rab8a (1 µM) in kinase assay buffer (25 mM Tris-HCl pH 7.4, 50 mM NaCl, 10 mM MgCl2, 0.5 mM GTP, 0.1 mg/ml BSA and 1 mM DTT). After addition of ATP (2 µM), kinase reactions were allowed to proceed for 10 or 30 minutes shaking at 650 rpm at 30 °C. No ATP was added to the control reaction which was allowed to proceed for 30 minutes shaking at 650 rpm at 30 °C. Kinase reactions were halted by adding SDS-PAGE loading buffer and reducing agent (1X, Invitrogen, catalog #NP0007 and NP0009) and boiling at 95 °C for 6 minutes. Samples were then loaded onto a 4-12% Bis-Tris gel (Invitrogen). Samples were immunoblotted as described above. Briefly, proteins were transferred onto PVDF membrane (Thermo Scientific, catalog #88520) at 200 mA for 2.5 hours at 4 °C. Following transfer and blocking in 5% milk or BSA for 1 hour at room temperature, membranes were cut above the 70 kDa marker (Chameleon Duo prestained protein ladder; LiCORbio, catalog #928-60000). The top portion of the membrane was immunoblotted for the RCKW portion of LRRK2 (N241A/34, RRID: AB_2877351) in 1% BSA, while the lower half was blotted for phospho-Rab8a T72 (MJF-R20, RRID: AB_2814988) and 6x-His (RRID: AB_3086567) in 1% milk overnight. Antibodies raised in different species were multiplexed as indicated in the text and figure legends. Blots were then washed three times for 10 mins with Tris-buffered saline supplemented with 0.1% Tween-20 (TBS-T), then incubated for 1 hour at room temperature with secondary antibodies (IRDye 680RD Goat Anti-Rabbit, RRID: AB_2721181 and IRDye 800CW Donkey Anti-Mouse, RRID: AB_2716622). Blots were washed a further three times in TBS-T and imaged using the LI-COR imaging system. Intensities of probed protein bands were measured using ImageJ (version 2.14.0). Statistical tests were performed as indicated in the text and figure legends using GraphPad Prism (version 10.5.0, RRID: SCR_002798).

### Figure preparation

Figures were prepared in Adobe Illustrator.

**Supplementary Figure 1.**
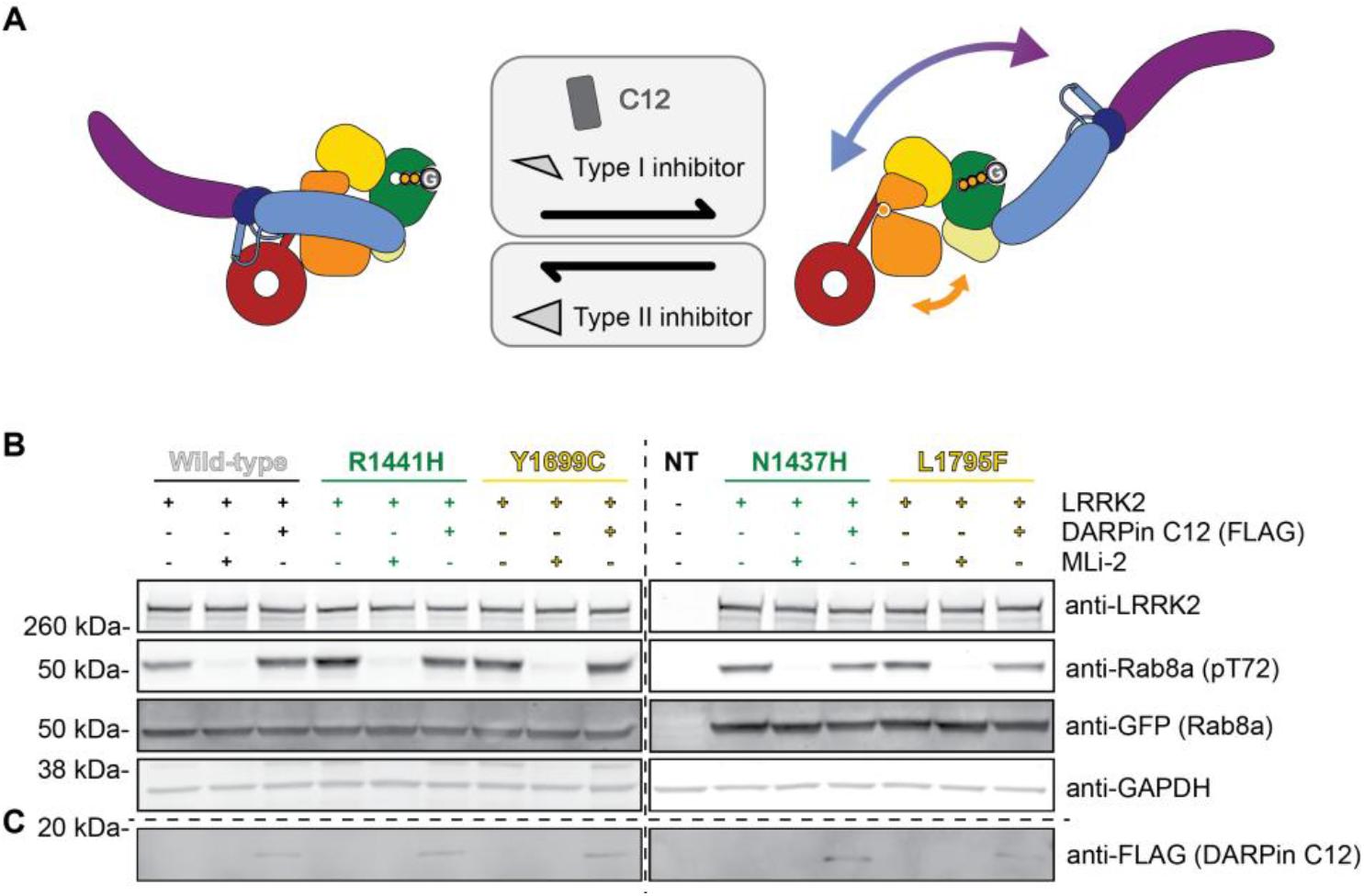
DARPin C12 and PD-associated LRRK2 ROC:COR-B mutants act at the same step of LRRK2 activation. **(A)** Schematic depiction of the conformational preference of different tools used in this study. DARPin C12 and type I kinase inhibitors such as MLi-2 promote the unlatched conformation of LRRK2. Type II kinase inhibitors such as RN277 promote the autoinhibited/latched conformation of LRRK2. **(B)** and **(C)** Representative immunoblots for Figure 1E. 293T cells overexpressing GFP-tagged Rab8a and various GFP-11-tagged LRRK2 variants co-expressed with (+) or without (-) FLAG-tagged DARPin C12 were treated with either DMSO (control) or MLi-2 (5 μM, 1 hour) before lysis and immunoblotting for LRRK2, GFP, Rab8a (pT72) and GAPDH (B) or FLAG (C). Samples were loaded identically and run on either Bis-Tris (B) or Tricine (C) gels under identical conditions (separate blots depicted by dashed lines). NT= no transfection. Blots were processed in ImageJ.

**Supplementary Figure 2.**
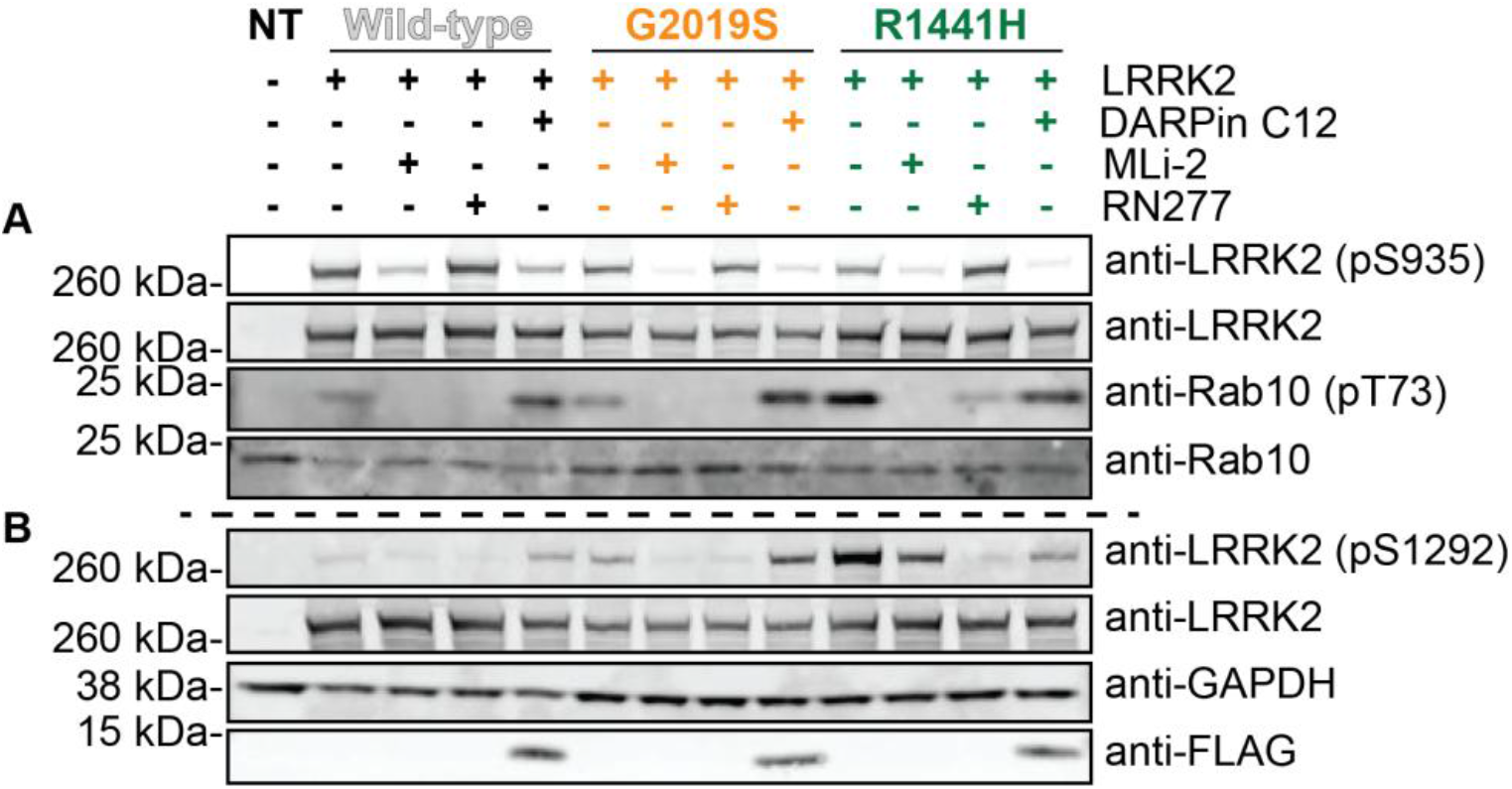
Representative immunoblots for Figure 2B and C. 293T cells overexpressing GFP11-tagged LRRK2 variants co-expressed with (+) or without (-) FLAG-tagged DARPin C12 were treated with either DMSO (control), MLi-2 (1uM), or RN277 (10uM) for 3 hours before lysis. Samples were loaded and run twice under identical conditions on Bis-Tris gels (separate blots depicted by dashed line) before immunoblotting for LRRK2 (pS935), LRRK2, Rab10 (pT73) and Rab10 (A) and LRRK2 (pS1292), LRRK2, GAPDH and FLAG (B). NT= no transfection. Blots were processed in ImageJ.

**Supplementary Figure 3.**
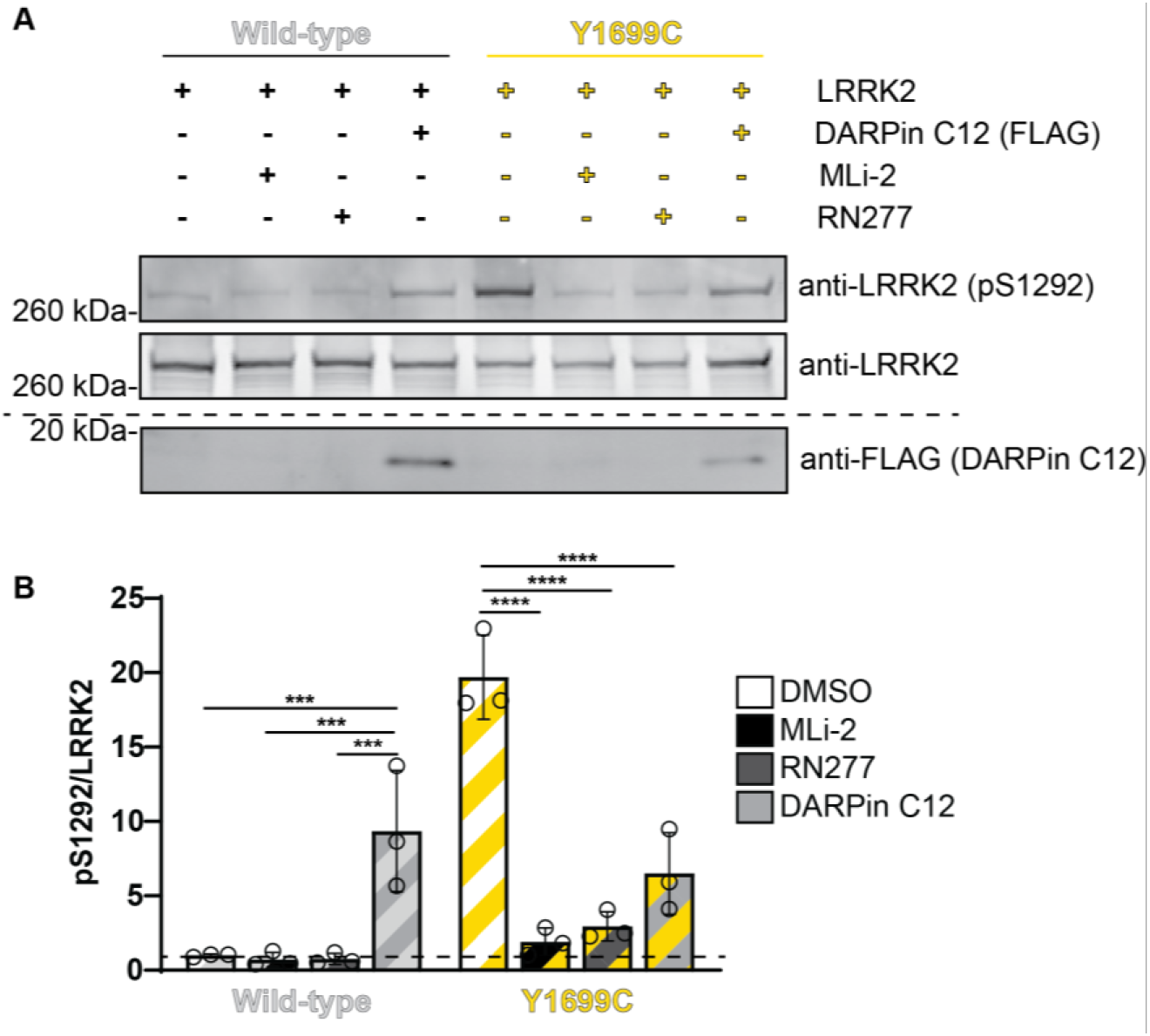
LRRK2[Y1699C] autophosphorylation profile mirrors LRRK2[R1441H]. **(A)** 293T cells overexpressing GFP11-tagged LRRK2[WT] or LRRK2[Y1699C] co-expressed with (+) or without (-) FLAG-tagged DARPin C12 were treated with DMSO (control), MLi-2 (1 μM) or RN277 (10 μM) for 3 hours before lysis. Samples were run on Bis-Tris gels and immunoblotted for LRRK2 (pS1292), and LRRK2. Samples were reprocessed and run on a Tricine gel to probe for FLAG (separate blots depicted by the dashed line). Blots were processed in ImageJ. **(B)** Quantification of LRRK2 autophosphorylation (LRRK2 (pS1292)/LRRK2, normalized to LRRK2[WT] DMSO condition) based on three (n=3) independent western blots (as shown in A). Blots were processed on ImageJ and data were analyzed using a two-way ANOVA with Tukey’s multiple comparison correction. *** p≤0.001; ****p≤0.0001; ns=non-significant. Error bars +/- SD.

**Supplementary Figure 4.**
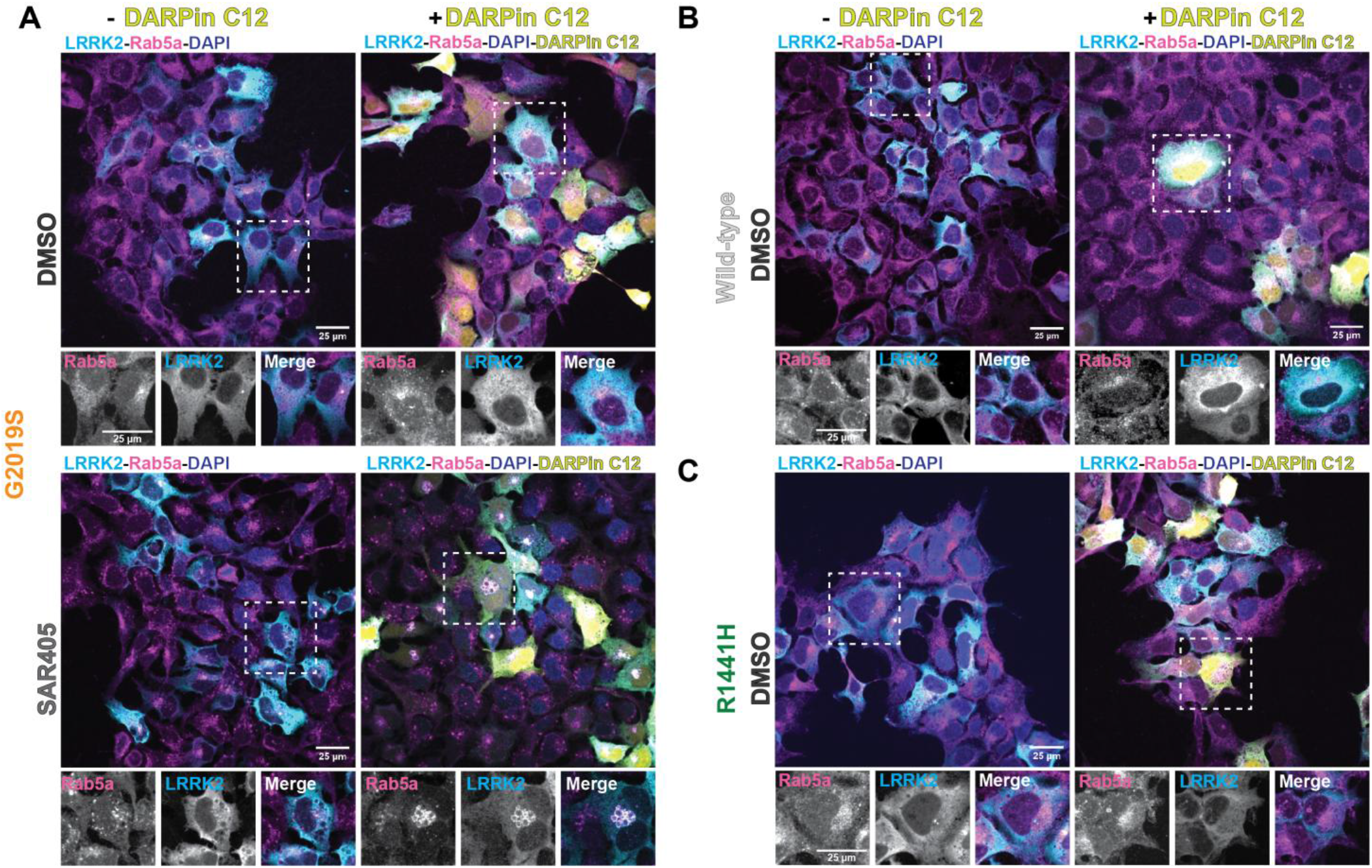
Additional representative images for Figure 3. Representative images of 293Ts overexpressing GFP11-tagged LRRK2[G2019S] (A), LRRK2[WT] (B) or LRRK2[R1441H] (C) without (-) or with (+) GFP-tagged DARPin C12 and immunostained for LRRK2, Rab5a, GFP-DARPin C12 and nuclei visualized with DAPI. Cells were either treated with DMSO (control) or the VSP34 inhibitor SAR405 (6μM, 30 minutes). Scale bars 25 μm.

**Supplementary Figure 5.**
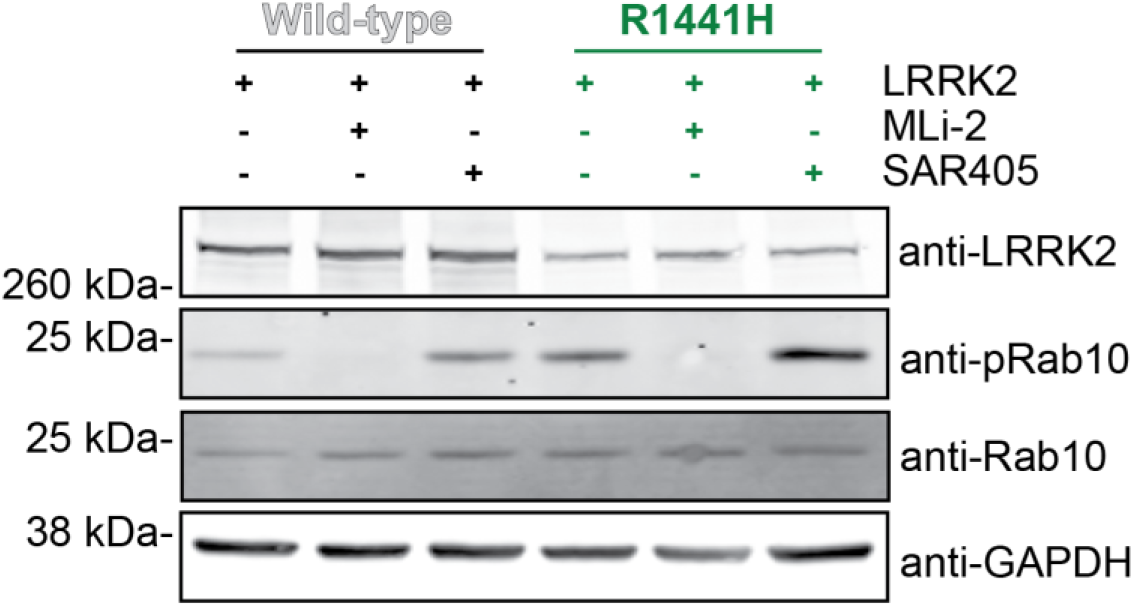
SAR405 stimulates increased substrate phosphorylation by LRRK2[WT] and LRRK2[R1441H]. 293Ts overexpressing either GFP11-tagged LRRK2[WT] or LRRK2[R1441H] were treated with either MLi-2 (5 μM; 1 hour), SAR405 (6 μM; 30 minutes) or DMSO before lysis. Samples were run on Bis-Tris gels and immunoblotted for LRRK2, Rab10 (pT73), Rab10, and GAPDH. Data representative of three independent experiments.

**Supplementary Figure 6.**
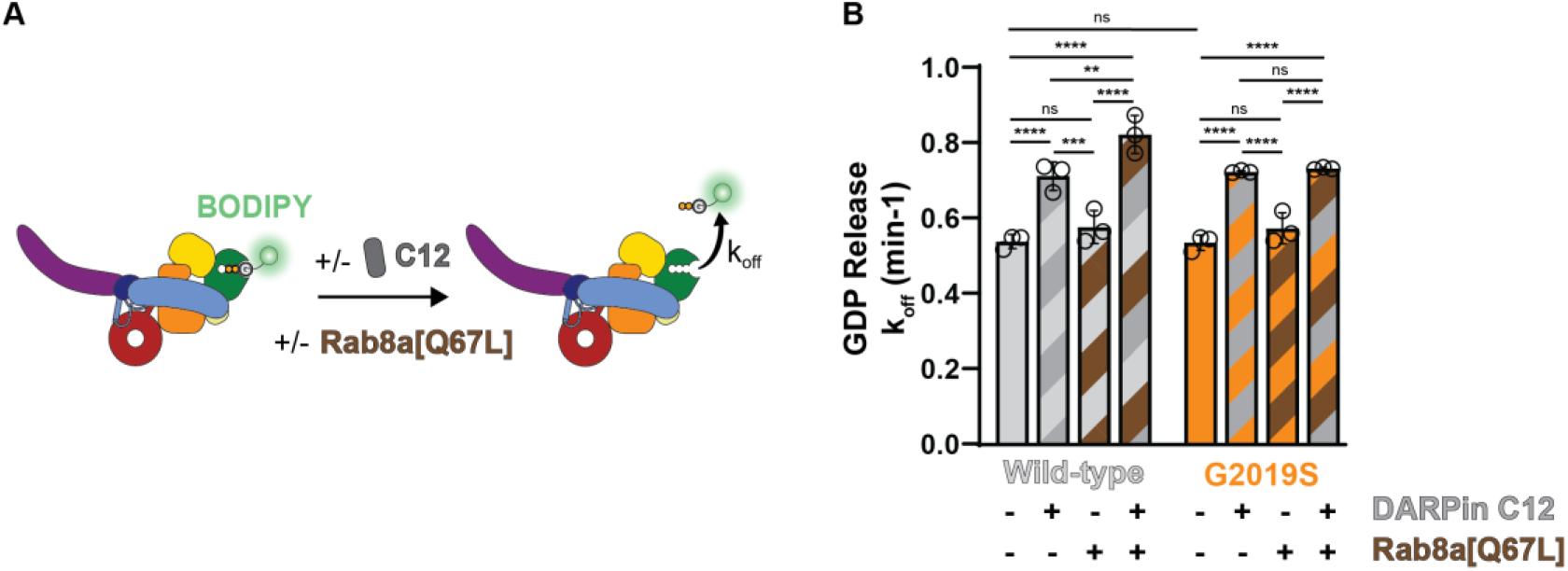
LRRK2[WT] and LRRK2[G2019S] both release GDP at a faster rate in the presence of DARPin C12. **(A)** The rate of BODIPY-FL-GDP release from LRRK2 *in vitro* was measured using a fluorescence-based assay in the presence (+) or absence (-) of DARPin C12 and Rab8a[Q67L]. **(B)** LRRK2[WT] and LRRK2[G2019S] both release GDP at a faster rate in the presence of DARPin C12. There is no significant difference (ns) in GDP release in the presence of Rab8a[Q67L] alone. Data from 3 independent experiments were analyzed using a two-way ANOVA with Tukey’s correction for multiple comparisons. ** p≤0.01; *** p≤0.001; **** p≤0.0001; ns=non-significant. Error bars ± SD.

**Supplementary Figure 7.**
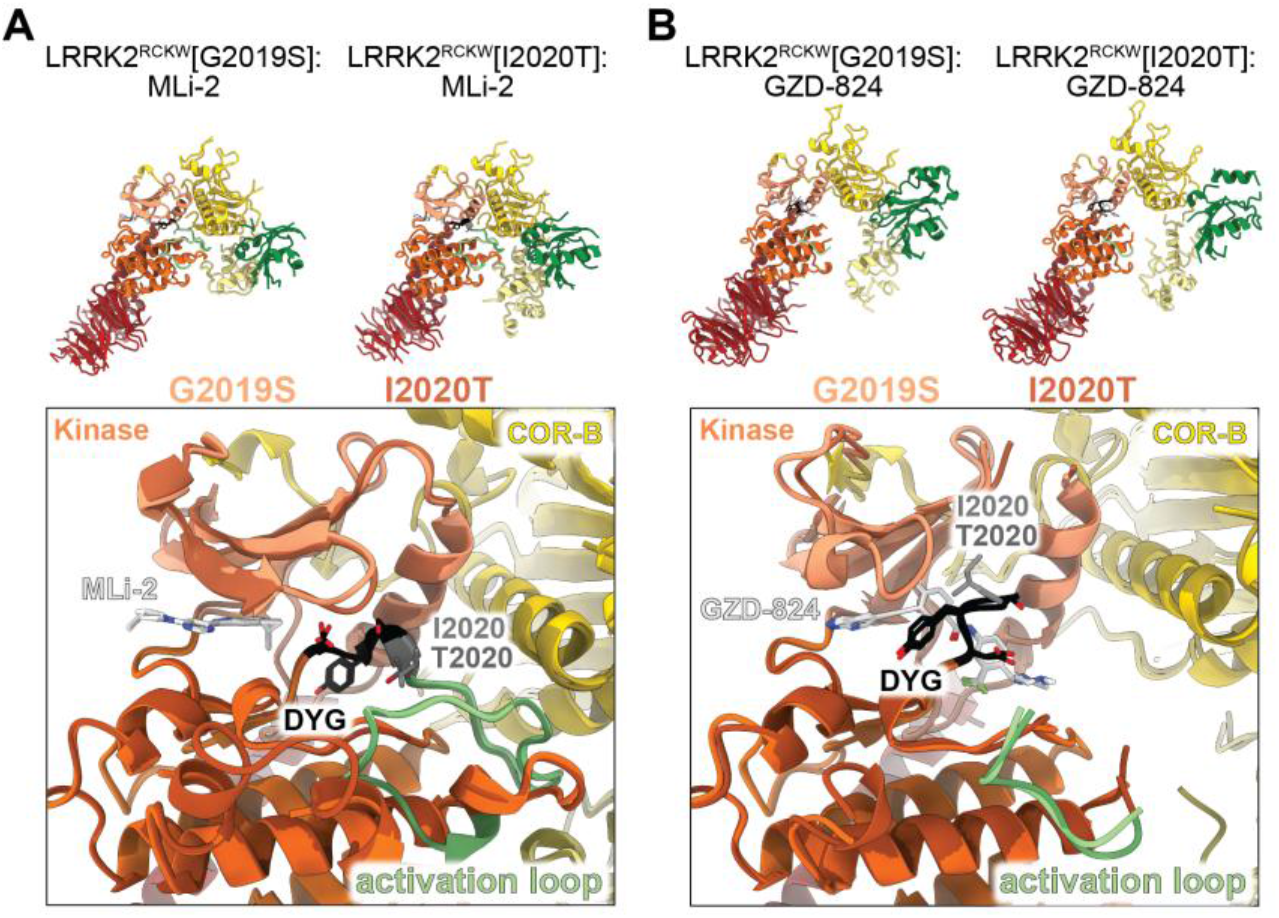
Comparison of RCKW[G2019S] and RCKW[I2020T] structures bound to inhibitors. **(A)** and **(B)** Comparisons of RCKW[G2019S] and RCKW[I2020T] active site bound to type I inhibitor MLi-2 (A) (LRRK2[G2019S] PDB: 8TZC; LRRK2[I2020T] PDB: 8TZG) or type II inhibitor GZD-824 (B) (LRRK2[G2019S] PDB: 8TYQ; LRRK2[I2020T] PDB: 8TZB)^50^. RCKW[G2019S] is portrayed by the lighter palette across all domains, while I2020T is portrayed in a darker palette. There is high overlap between RCKW[G2019S] and RCKW[I2020T], indicating no structural reason for differences in LRRK2[G2019S] and LRRK2[I2020T] behaviors.

**Supplementary Figure 8.**
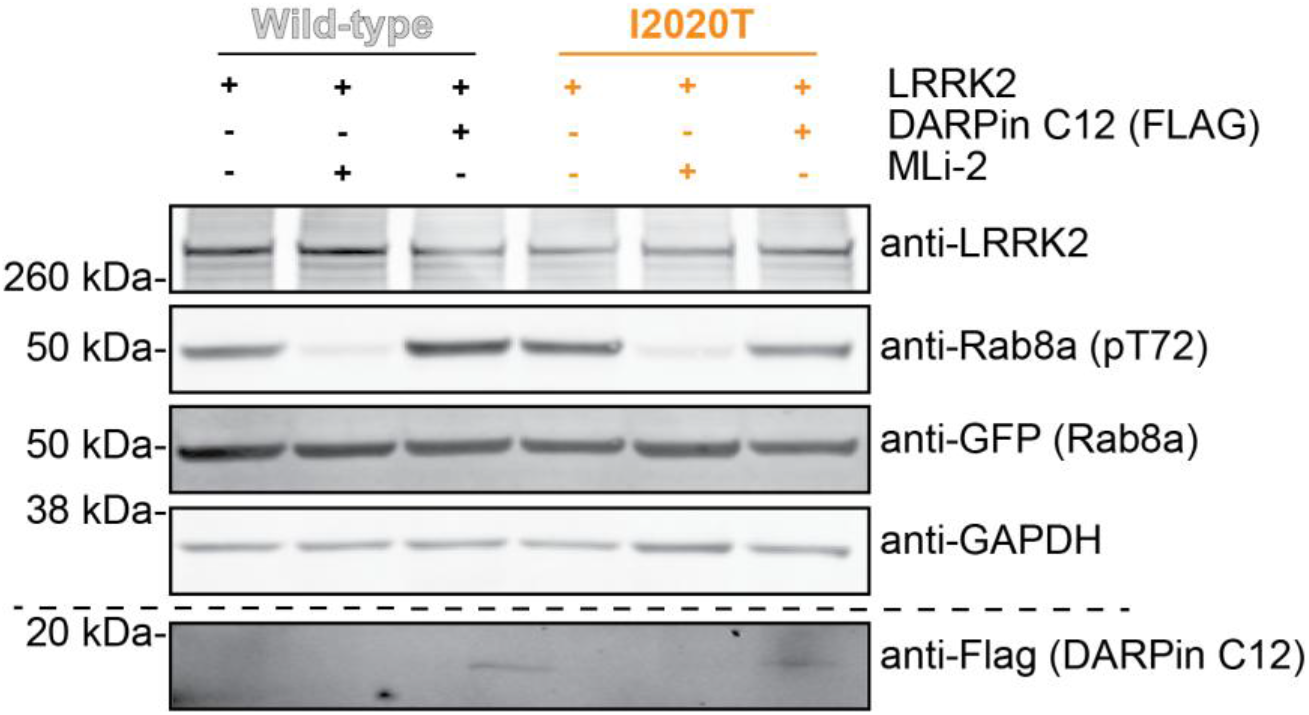
Representative immunoblots for Figure 5A. 293T cells expressing GFP-tagged Rab8a, either GFP11-tagged LRRK2[WT] or [I2020T] and co-transfected with (+) or without (-) FLAG-tagged DARPin C12 were treated with DMSO or MLi-2 (1 μM) for 1 hour before lysis. Samples were run on a Bis-Tris gel and immunoblotted for LRRK2, Rab8a (pT72), GFP, and GAPDH. Samples were reprocessed and run on a Tricine gel to probe for FLAG (separate blots depicted by dashed line). Blots were processed in ImageJ.

**Supplementary Figure 9.**
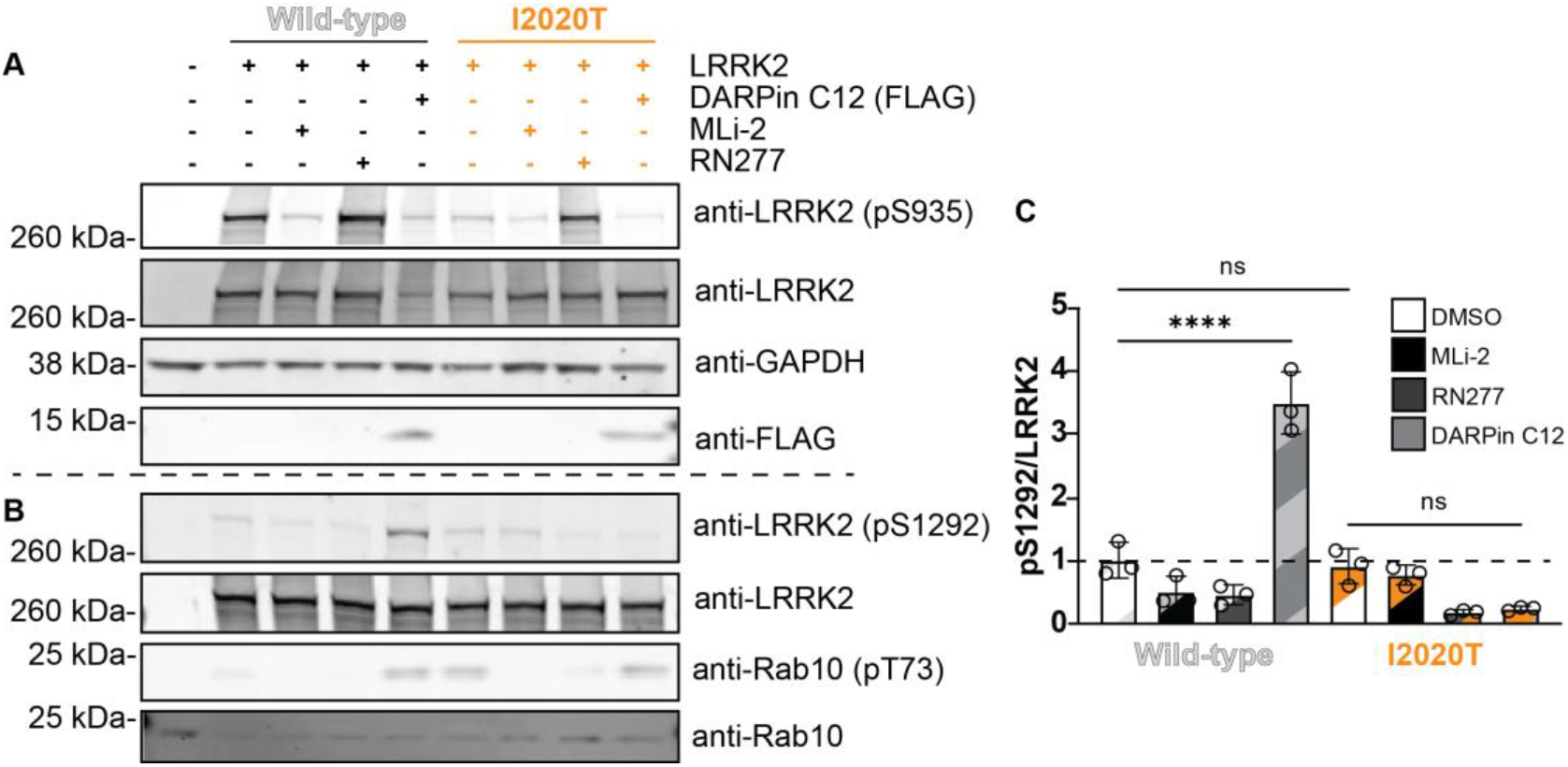
LRRK2[I2020T] has reduced autophosphorylation in the presence of DARPin C12. **(A, B)** Representative immunoblots for Figure 5B. 293T cells overexpressing GFP11-tagged LRRK2[WT] or LRRK2[I2020T] co-expressed with (+) or without (-) FLAG-tagged DARPin C12 were treated with either DMSO, MLi-2 (1uM), or RN277 (10uM) for 3 hours before lysis and immunoblotting. Samples were run twice under identical conditions (separate blots depicted by dashed line) and immunoblotted for LRRK2, LRRK2 (pS935), GAPDH and FLAG and LRRK2 (A) or LRRK2 (pS1292), LRRK2, Rab10 (pT73) and Rab10 (B). Blots were processed in ImageJ. **(C)** Quantification of LRRK2[WT] and LRRK2[I2020T] autophosphorylation at residue S1292 when treated with DMSO, MLi-2 (1 μM), or RN277 (10 μM) for 3 hours, or in the presence of FLAG-tagged DARPin C12 from 3 independent western blots (n=3) **** p≤0.0001; ns=non-significant. Error bars ± SD. Representative immunoblot related to this analysis is shown in B.

