## Supplementary Table 1 for "Conformation governs cellular behaviors of LRRK2 Parkinson’s disease variants"

| ANTIBODY NAME | WORKING CONCENTRATION | SOURCE | IDENTIFIER |
|---|---|---|---|
| LRRK2 (MJFF2-c41-2) | 1:1000, WB | Abcam | Cat# ab133474; Lot# GR3445288–3; RRID: AB_2713963 |
| LRRK2/Dardarin (N241A/34) | 1:1000, WB | NeuroMab (UC Davis) | Cat# 75-253-020; Lot# 455.7JD.23H; RRID: AB_2877351 |
| LRRK2 [UDD3 30(12)] | 1:200, IF | Abcam | Cat# ab133518; Lot# 1128968-4; RRID: AB_2938806 |
| Phospho-LRRK2 (pS1292; MJFR-19-7-8) | 1:1000, WB | Abcam | Cat# ab203181; Lot# GR3259813-8; RRID: AB_2921223 |
| Phospho-LRRK2 (pS935; UDD2 10(12)) | 1:1000, WB | Abcam | Cat# ab133450; Lot# 1003146-74; RRID: AB_2904231 |
| GFP | 1:2000, WB | Santa Cruz Biotechnology | Cat# sc-9996; Lot# C2523; RRID: AB_627695 |
| 6x-His | 1:2000, WB | Proteintech | Cat# 66005; Lot# 10027680; RRID: AB_3086567 |
| Phospho-Rab8a (pT72; MJF-R20) | 1:1000, WB | Abcam | Cat# ab230260, Lot# GR3216587–1; RRID: AB_2814988 |
| Phospho-Rab10 (pT73; MJF-R21) | 1:500, WB | Abcam | Cat# ab230261; Lot# GR3216588-1; RRID: AB_2811274 |
| Rab10 (605B11) | 1:500, WB | Nanotools | Cat# 0680-100; Lot# 068050201; RRID: AB_2921226 |
| GAPDH | 1:8000, WB | Cell Signaling Technology | Cat# 2118; Lot# 16; RRID: AB_561053 |
| FLAG | 1:500, WB | PGT Labs | Cat# 20543-1-AP, Lot# 00106090; RRID: AB_11232216 |
| Rab5a | 1:200, IF | Invitrogen | Cat# 14-9711-82, Lot# 7; RRID: AB_2884043 |
| IRDye 680RD Goat Anti-Rabbit | 1:10,000, IF | LI-COR Biosciences | Cat# 926-68071, Lot# D50729-04; RRID: AB_2721181 |
| IRDye 800CW Donkey Anti-Mouse | 1:10,000, IF | LI-COR Biosciences | Cat# 926-32212; Lot# D40820-06; RRID: AB_2716622 |
| Alexa Fluor 586 Goat Anti-Mouse IgG (H+L) | 1:200, IF | Invitrogen | Cat #A11004, Lot #2198584 |
| Alexa Fluor 647 Chicken Anti-Rabbit IgG (H+L) | 1:200, IF | Life Technologies | Cat #A21443, Lot # 1784026 |
